# Western-style diet-induced microbial shifts and inflammation reprogram *c-Kit*⁺ secretory cells into targets for tumor initiation

**DOI:** 10.64898/2026.04.06.716383

**Authors:** Sara Silva, Paola Procopio, Valeriya Zinina, Solveig Turß, Rafael R Segura Munoz, Seyram Segbefia, Sena Bae, Lior Lobel, Luigi Nardella, Andrea Sacchetti, Rosalie Joosten, Coen Linn, Susanna Plugge, Mathijs P Verhagen, Fatma Aktuna, Kim Pauck, Nicole Löwer, Victor Sourjik, Holger Garn, Richard S. Blumberg, Jens Puschhof, Wendy S. Garrett, Riccardo Fodde, Mark Schmitt

**Author notes:** to whom correspondence should be addressed at: Dr. Mark Schmitt, Institute of Pharmacology, University of Marburg, Address: Karl-von-Frisch-Straße 2, 35043 Marburg, Germany.

## Abstract

While colorectal cancer (CRC) is thought to originate primarily from resident stem cells, diet heavily influences intestinal stem cell homeostasis and disease onset. In this study, we demonstrate that a Western-style diet (WSD) alters intestinal homeostasis by uncoupling canonical stemness from tumorigenesis. WSD exposure suppresses canonical *Lgr5*⁺ stem cells while activating an alternative pool of facultative *c-Kit*⁺ secretory cells. Although these reprogrammed stem-like cells exhibit genotoxic stress, they remain proliferative under prolonged dietary exposure, suggesting increased susceptibility to tumor-initiating mutations. We identify diet-induced microbial shifts and the expansion of enterotoxigenic *Bacteroides fragilis* (ETBF) as the upstream driver. ETBF and its toxin fragilysin autonomously trigger multipotency in *c-Kit*^+^ cells through Wnt/β-catenin and YAP signaling. Importantly, these alterations are fully reversible upon dietary intervention. Together, our results highlight a targetable dietary-microbial axis that shapes epithelial stemness and underscores the role of facultative stem cells in diet-induced CRC onset.

## Introduction

One of the main dogmas in cancer biology predicts that most malignancies arise from resident stem cells. However, as we recently demonstrated in mouse models and validated in patient-derived colon cancers, in the context of inflammation, intestinal adenomas rather originate from committed lineages. This observation is of relevance for the so-called “bad luck cancer” debate, triggered by the Tomasetti & Vogelstein study^1^, according to which cancer risk is mainly determined by the rate at which stem cells divide and accumulate spontaneously occurring somatic mutations. However, in their mathematical model, stem cell division and environmental factors were regarded as two independent variables, somehow implying that etiologic risk factors such as diet and inflammation in colon cancer do not affect the rate at which stem cells divide^2^. This is in contradiction with our current knowledge of the intestinal epithelium as one of the most rapidly renewing tissues in mammals maintained, in homeostasis, by a pool of cycling Leucine-rich repeat-containing G-protein coupled receptor 5 expressing intestinal stem cells (*Lgr5*⁺ ISCs), located at the base of the crypts^3^. However, in response to tissue damage, such as chemical- or radiation-induced tissue injury, the intestinal epithelium exhibits remarkable plasticity, whereby differentiated or lineage-committed cells can revert to a stem-like state to support intestinal regeneration. This facultative stem cell activity has been described in multiple intestinal lineages, including Paneth cells (PCs) in the small intestine (SI) and deep crypt secretory (DCS) cells in the colon, particularly in the context of severe inflammation and/or tissue damage^4–8^.

While beneficial for the tissue’s regenerative response, the recruitment of lineage-committed intestinal epithelial cells (IECs) into the stem cell pool may be deleterious, as their reprogramming can confer tumor-initiating capacity, as shown in experimental mouse models of acute dextran sodium sulphate (DSS)-driven) inflammation and in patients with a history of inflammatory bowel diseases (IBD)^9–12^. Of note, we observed that colon cancer initiation from committed cells is not necessarily restricted to patients suffering from IBD, but also holds true for a substantial proportion of sporadic, non-IBD cases^12^. As such, it is plausible to hypothesize that increased exposure to environmental risk factors leading to subclinical but persistent low-grade inflammation may exert comparable reprogramming effects.

From this perspective, Western-style diet (WSD), a main modifiable colon cancer risk factor, characterized by high fat and low fibre content and strongly associated with colon cancer^13^, is shown to activate myeloid cells and induce innate immune responses leading to a subclinical, though persistent type of metabolic inflammation, termed metaflammation^14,15^. Here, we investigated whether the WSD-driven inflammation alters the composition of the intestinal stem cell niche and expands the pool of cell targets for tumor initiation, similar to what was observed upon acute inflammation in mice (DSS-driven) and men (IBD)^4,11^.

We found that WSD rapidly suppresses canonical *Lgr5*⁺ ISCs while enhancing epithelial stemness through activation of reserve stem cell programmes in small intestinal PCs and DCS cells of the colon. The reprogrammed PCs and DCS cells undergo genotoxic and inflammatory stress, rendering them vulnerable to somatic mutations and positioning them as potential drivers of tumorigenesis. The inflammation-associated reprogramming is driven, at least in part, by alterations of the microbiome, including expansion of enterotoxigenic *Bacteroides fragilis (B. fragilis)*.

Collectively, our findings reveal that dietary cues can reversibly rewire intestinal stem cell identity through inflammation and microbiota-dependent mechanisms, providing a functional link among diet, the gut microbiome, and epithelial plasticity.

## Results

### A rodent Western-style diet induces low-grade intestinal inflammation and hyperproliferation in the mouse intestine

To investigate the consequences of Western-style dietary habits on the intestinal tract *in vivo*, we employed the New Western-Style Diet 1 (NWD1), a purified AIN76A-based rodent diet that reflects dietary patterns associated with increased colon cancer risk in humans^16^.

First, we fed 8-week-old mice that had been maintained on the AIN76A-control diet from weaning with the NWD1 diet for one month and analyzed their intestinal tissues for proinflammatory changes. Despite the absence of macroscopically visible effects (Supplementary Fig. S1A), immunofluorescence and RT-qPCR analyses revealed increased numbers of total and tissue infiltrating CD3^+^ T cells, CD11^+^ dendritic cells and Ly6G^+^ neutrophils, as well as elevated TNFα, IL-1a/b, and IL-10 levels in NWD1-fed mice compared to control mice that remained on the AIN76A diet (Fig. 1A-E and Supplementary Data Fig. 1B). Together these findings indicate that exposure to rodent Western-style diet is sufficient to induce a low-grade inflammatory response in the gut. Furthermore, we observed an overall expansion of Ki67^+^ cells within small intestinal (SI) crypts of NWD1-fed mice (Fig. 1F). This was most evident at the crypt base, where continuous nuclear Ki67 saining was detected throughout the entire cell compartment, in contrast to crypts from AIN76A-fed mice, which displayed the characteristic alternating pattern of Ki67^+^ *Lgr5*⁺ ISCs and Ki67^−^ PCs (Fig. 1G). A comparable increase in proliferating cells was also observed in colonic crypts (Supplementary Fig. 1C). These findings demonstrate that, in addition to eliciting a proinflammatory response, the NWD1 diet promotes epithelial proliferation throughout the intestine.

**Figure 1:**
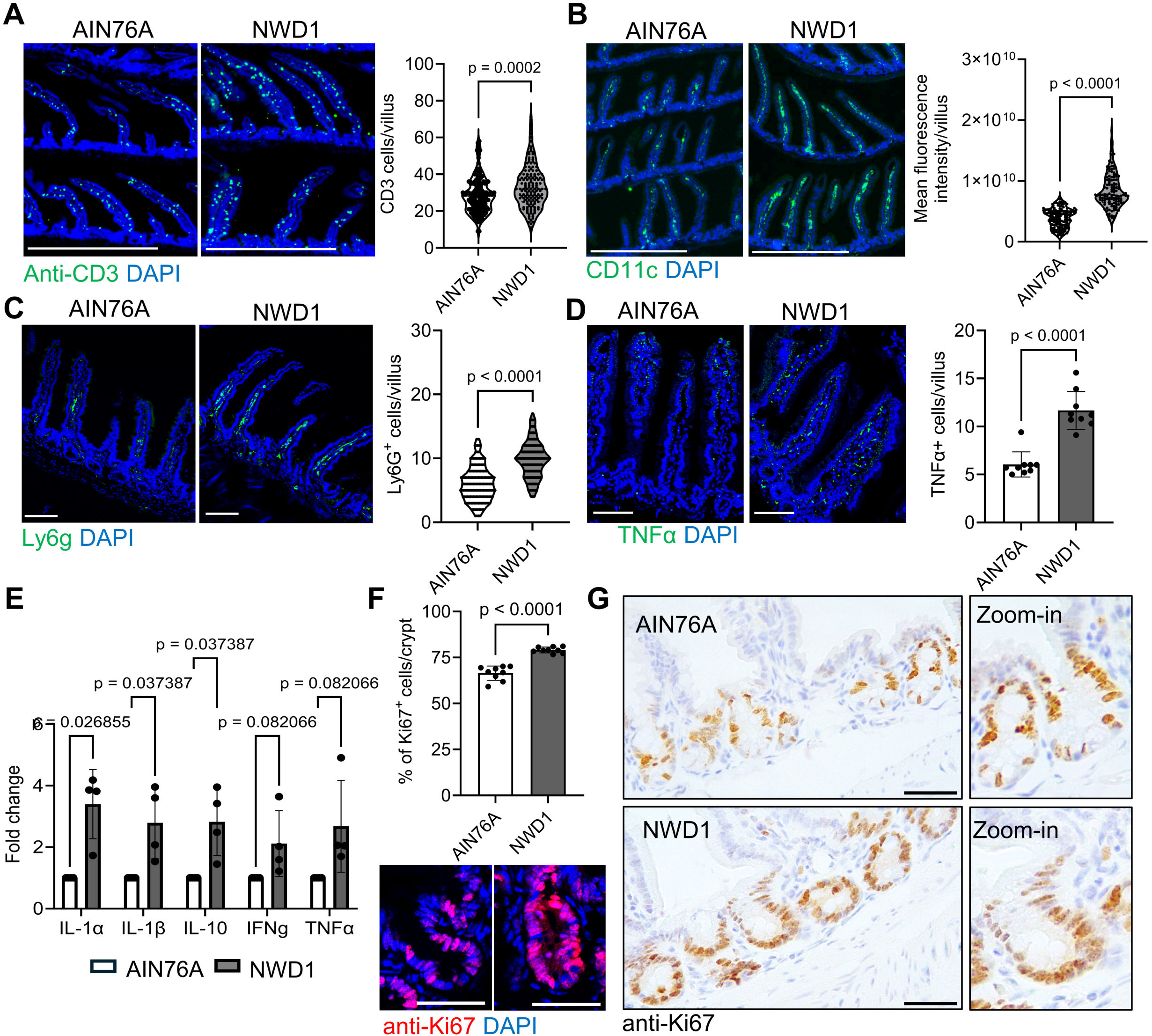
NWD1 induces inflammation and hyperproliferation in the intestine. **A)** Confocal microscopy analysis and quantification of CD3^+^ cells in small intestinal tissues of mice fed for 1 month with AIN76A or NWD1. Graph shows the quantification of total CD3^+^ cell numbers/villus (results represent pooled data of 3 individual mice per group, scale bars = 500 µm, nuclei were counterstained with DAPI). **B)** Analysis of CD11c expression in 1-month AIN76A and NWD1 fed mice (scale bars = 500um, nuclei were counterstained with DAPI). Graph shows mean intensity of CD11c fluorescence/villus (results represent pooled data of 3 individual mice per group). **C)** Analysis and quantification of Ly6G^+^ cells in 1-month AIN76A and NWD1 fed mice. Graph shows the quantification of total Ly6G^+^ cell numbers/villus (results represent pooled data of 3 individual mice per group, scale bars = 100 µm, nuclei were counterstained with DAPI). **D)** Analysis of TNFα expression in 1-month AIN76A and NWD1 fed mice (scale bars =100um, nuclei were counterstained with DAPI). Graph shows the quantification of total TNFα^+^ cell numbers/villus (results represent pooled data of 3 individual mice per group). **E)** RT–qPCR analysis of the indicated genes in intestinal crypts from 1-month AIN76A or NWD1 fed mice (n = 4 biological replicates). **F)** Confocal microscopy analysis and quantification of Ki67^+^ cells in SI crypts of 1-month AIN76A and NWD1 fed mice (results represent pooled data of 3 individual mice per group, scale bar= 50 µm, nuclei were counterstained with DAPI). **G)** Ki67 IHC analysis of mouse small intestinal tissues of 1-month AIN76A and NWD1 fed mice (n= 3 mice per group, scale bar= 75 µm). All data are mean ± s.d. and were analyzed by two-tailed Student’s t-test (**A–F**).

### NWD1 suppresses canonical *Lgr5*^+^ ISCs while increasing intestinal proliferation and stemness of secretory Paneth and DCS cell lineages

To address how the pro-inflammatory NWD1 influences the proliferative capacity of specific cell types located in the ISC niche, we fed the NWD1 and the AIN76A diets to genetically modified mice, allowing the monitoring and lineage tracing of three crypt-resident intestinal lineages: *Lgr5*^EGFP-IRES-CreERT2^;*Rosa*^LSL-dTomato^ mice, tagging *Lgr5⁺* ISCs in the SI and colon^3^; *Lyz*^CreERT2^/*Rosa*^LSL-YFP^ mice, marking mature PCs in the SI^5,12^; and *c-Kit*^CreERT2^; *Rosa*^LSL-dtomato^ mice, labelling PCs in the SI and DCS in the colon^17^.

Consistent with the results from previous studies reporting on the WSD-mediated inhibition of canonical ISCs in the SI^18,19^, we observed significantly reduced numbers of *Lgr5*⁺ ISCs in both the SI and colon of 3-months NWD1-fed mice (Supplementary Fig. S2A and B). More importantly, the NWD1 diet nearly completely suppressed the lineage tracing and proliferative capacity of the remaining *Lgr5*⁺ cells (Fig. 2A and B).

**Figure 2:**
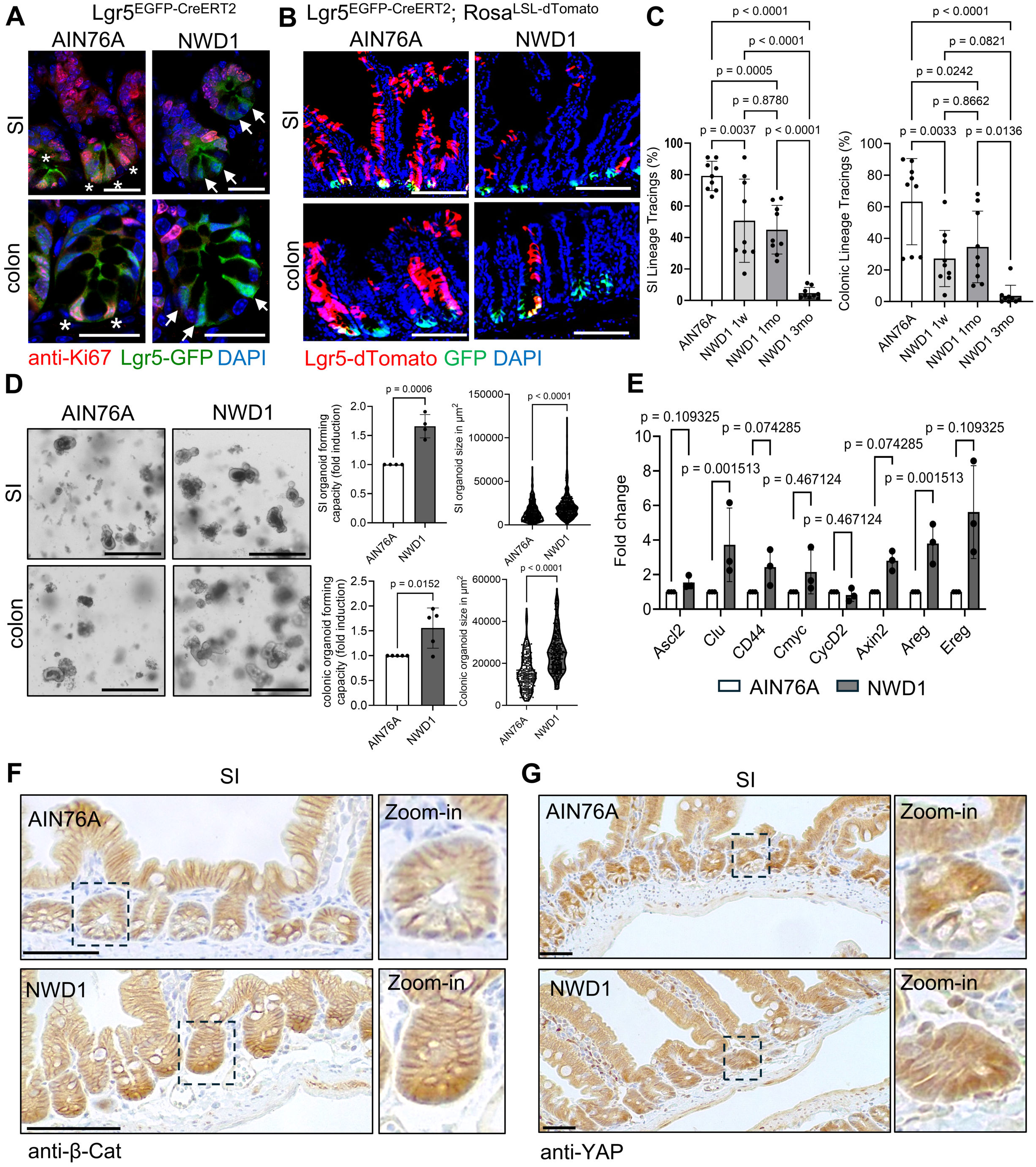
NWD1 suppresses *Lgr5*^+^ ISCs but increases intestinal stemness. **A)** Confocal microscopy analysis of intestinal tissues of *Lgr5*^EGFP-IRES-CreERT2^ mice fed with AIN76A or NWD1 for 3 months using antibodies against Ki67. Asterisks indicate Ki67^+^/GFP^+^ *Lgr5*^+^ ISCs in crypts of AIN76A fed mice, arrows indicate Ki67^-^/GFP^+^ *Lgr5*^+^ ISCs in crypts of NWD1 fed mice (n=3; nuclei were counterstained with DAPI). **B)** Confocal microscopy images of intestinal tissues of *Lgr5*^EGFP-IRES-creERT2^/*R26*^LSL-dTomato^ mice fed for 3 months with AIN76A or NWD1 (n = 3 mice per group; scale bar = 100 µm; nuclei were counterstained with DAPI). **C)** Quantification of lineage tracing events in *Lgr5*^EGFP-IRES-creERT2^/*R26*^LSL-dTomato^ fed with AIN76a and NWD1 diets for the indicated timepoints (results represent pooled data of 3 individual mice per group) **D)** Organoid multiplicities and sizes from SI and colonic crypts of 1-month AIN76A and NWD1 fed mice. (n= 6 for SI and 5 for colonic organoids, scale bars = 200 µm). **E)** RT–qPCR analysis of the indicated genes in intestinal crypts from 1-months AIN76A or NWD1 fed mice (n = 4 biological replicates). **F)** β-Catenin IHC analysis of SI tissues of mice fed for 1 month with AIN76A or NWD1 (n=3 per group, scale bar = 75 µm, box marks the area of higher magnification images). **G)** YAP IHC analysis of SI tissues of mice fed for 1 month with AIN76A or NWD1 (n=3 mice per group, scale bar =75 µm, box marks the area of higher magnification image). All data are mean ± s.d. and were analyzed by one-way analysis of variance (ANOVA) (**C**) with Tukey’s multiple comparison test or two-tailed Student’s *t*-test (**D E**).

Next, to determine the onset of this stem cell–suppressive effect, we progressively shortened the duration of NWD1 feeding and found that the lineage-tracing capacity of *Lgr5*⁺ ISCs was significantly impaired after just 1 week on the experimental diet (Fig. 2C). Unexpectedly, despite this suppression of canonical stem cell activity, both the multiplicity and size of organoids derived from small intestinal and colonic crypts isolated from NWD1-fed mice were increased compared with those from AIN76A-fed controls (Fig. 2D). These findings suggest that, although the rodent WSD suppresses canonical ISCs, it enhances the overall stemness of the intestinal niche. Consistent with these findings, qRT-PCR analysis revealed upregulation of stemness-associated genes in small intestinal crypts, including Wnt and YAP target genes, as well as *Clusterin*, a marker of revival stem cells (RSCs) ^20^ (Fig. 2E). The increased YAP and β-catenin activation in small intestinal and colonic crypts from NWD1-fed mice was further confirmed by immunohistochemical analysis of intestinal tissues following 1 month of experimental diet exposure (Fig. 2F and G and Supplementary Fig. S2C and D).

The above results are highly reminiscent of our previous findings in acute, DSS-driven inflammation, where PCs were identified as a reserve ISC population contributing to tissue regeneration and to tumor onset^4,5^. To determine whether WSD, similar to acute inflammation, can induce dedifferentiation of secretory intestinal lineages into stem-like cells, we isolated PCs, as well as *Lgr5⁺*ISCs from AIN76A- and NWD1-fed mice by FACS and assessed their capacity to autonomous organoid-forming capacity *ex vivo*^21^. *Lgr5⁺* ISCs can form organoids only in the presence of PCs and, accordingly, failed to do so when plated as single cells, irrespective of diet. In contrast, as previously observed in PCs from DSS-treated mice^5^, FACS-isolated PCs from NWD1-fed mice autonomously formed organoids, indicative of their acquisition of stem–like properties (Supplementary Fig. 3A).

To validate *in vivo* the NWD1-driven reprogramming of PCs into stem-like cells, we employed *Lyz*^CreERT2^/*Rosa*^LSL-YFP5,12^ and *c-Kit*^CreERT2^/*Rosa*^LSL-dtomato^ mice^17^ to monitor and lineage-trace mature PCs and DCS cells in the small and large intestine, respectively. Mice fed with NWD1 for three months showed significantly increased lineage tracing from *Lyz*^+^ PCs when compared with AIN76A-fed control animals, thus demonstrating *in vivo* that the Western-style rodent diet reprograms PCs into stem-like cells in the absence of functional *Lgr5*^+^ ISCs (Fig. 3A).

**Figure 3:**
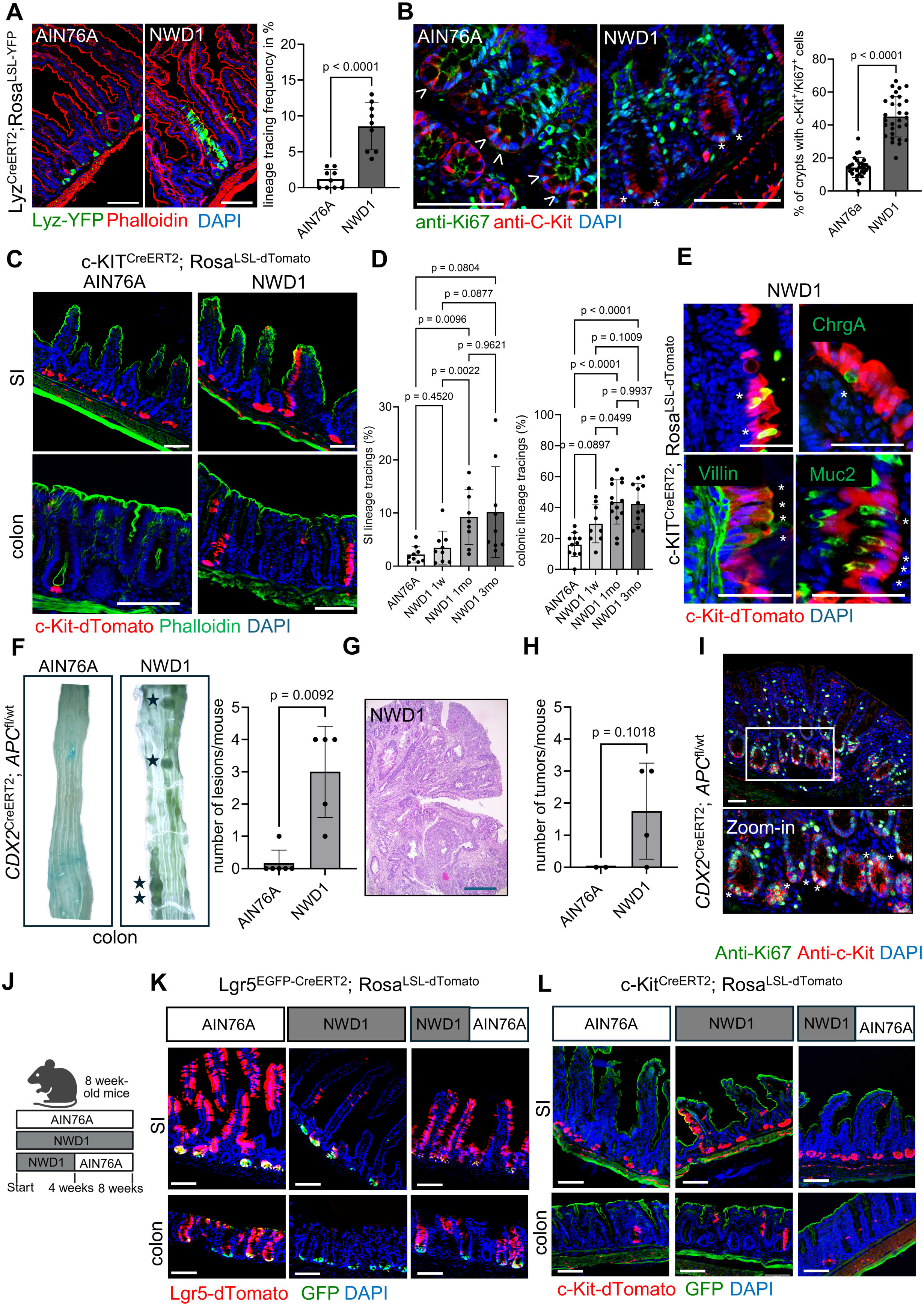
NWD1 reversibly reprograms small intestinal PCs and colonic DCS cells to stem-like cells and enhances intestinal tumorigenesis. **A)** Lineage tracing analysis of tamoxifen-injected *Lys*^CreERT2^/*R26*^LSL-YFP^ mice fed for 3 months with AIN76A or NWD1 diets (scale bar = 100 µM, tissues were counterstained with DAPI and phalloidin). The graph shows the quantification of the lineage tracing frequency of YFP labelled Paneth cells (results represent pooled data of 3 individual mice per group). **B)** Confocal microscopy images of c-Kit/Ki67 double positive cells, located at the crypt base in colonic tissues of mice fed for 1 month with AIN76A or NWD1 diets. Arrows indicate Ki67^+^/c-Kit^-^ *Lgr5*^+^ ISCs in AIN76A tissues, asterisks indicate Ki67^+^/c-Kit^+^ DCS cells in NWD1 tissues (scale bar = 100 µM, nuclei were counterstained with DAPI). The graph shows the quantification of Ki67^+^/c-Kit^+^ cells in mice fed with the indicated diets (results represent pooled data of 3 individual mice per group). **C)** Lineage tracing analysis of tamoxifen-injected *c-Kit*^CreERT2^/*R26*^LSL-dTomato^ mice fed for 1 month with AIN76A or NWD1 diets (scale bar = 100 µm, tissues were counterstained with phalloidin and DAPI). **D)** Quantification of the lineage tracing frequency of *c-Kit*^+^ PCs in the SI (left) and *c-Kit*^+^ DCS cells in the colon (right) of mice fed for the indicated timepoints with AIN76A or NWD1 diets (results represent pooled data of 3 individual mice per group). **E**) Analysis of Villin, Muc2, Dclk1 and Chromogranin A expression in 1-month NWD1 fed *c-Kit*^CreERT2^/*R26*^LSL-dTomato^ mice 10 days after tamoxifen treatment. Asterisks indicate double positive cells (n=3 mice per group, scale bar = 50µm, nuclei were counterstained with DAPI). **F)** Colonic tissues of *CDX2*^CreERT2^; *APC*^fl/wt^ fed for 6 months with AIN76A or NWD1 diets after tamoxifen induction. Tissues were stained with methylene-blue and visible polyps (asterisks) were quantified (n = 5 mice per group). **G)** Representative H&E staining of a colorectal adenocarcinoma of NWD1-fed *CDX2*^CreERT2^; *APC*^fl/wt^ mice, 9 months post tamoxifen induction (scale bar = 500 µm). **H)** Quantification of tumors of *CDX2*^CreERT2^; *APC*^fl/wt^ mice fed for 9 months with experimental diets (n = 2 for AIN76A and 4 for NWD1). **I)** Co-immunostaining using antibodies against c-Kit and Ki67 on colonic tissues of NWD1 fed *CDX2*^CreERT2^; *APC*^fl/wt^ mice. Box marks the area of magnification image. Asterisks indicate c-Kit/Ki67 double positive cells (n = 5 mice, scale bar = 50 µm). **J)** Feeding regimen of mice described in (**K** and **L**). Mice were either maintained on AIN76A, fed with NWD1 for 2 months or fed for 1 month with NWD and then switched back to AIN76A for 1 month. **K)** Lineage tracing analysis of *Lgr5*^EGFP-IRES-creERT2^/*R26*^LSL-dTomato^ mice treated according to the regimen described in (**J**) (n= 3 mice per group, scale bar= 100 µm, nuclei were counterstained with DAPI). **L)** Lineage tracing analysis of *c-Kit*^CreERT2^/*R26*^LSL-dTomato^ mice treated according to the regimen described in (**J**) (n= 3 mice per group, scale bar= 100 µm, tissues were counterstained with phalloidin and DAPI). All data are mean ± s.d. and were analyzed by two-tailed Student’s *t*-test (**A**, **B, F** and **H**) or one-way ANOVA (**D**) with Bonferroni’s multiple comparison test.

As PCs are exclusively resident in the SI, we next examined whether the WSD-like NWD1 had similar effects on DCS, the colonic equivalent of PCs, previously shown to acquire multipotency during acute colitis^22–24^. Similar to PCs, DCS cells can be identified by their specific location flanking *Lgr5*^+^ ISCs at the base of colonic crypts, as well as by the expression of the tyrosine kinase receptor *c-Kit*^22,24^. To determine if DCS cells acquire proliferative capacity in the context of WSD, we first examined colonic tissues from NWD1-fed mice by co-immunostaining with c-Kit and Ki67 antibodies. A significant increase in the number of double-positive cells at the crypt base was observed in NWD1-fed mice when compared with the animals fed with the AIN76A control diet, thus suggesting the WSD-driven activation of DCS cell proliferation (Fig. 3B).

Next, we addressed the lineage-tracing capacity of *c-Kit*^+^ cells and fed *c-Kit*^CreERT2^; *Rosa*^LSL-dtomato^ mice^17^ for 1 week to 3 months with the NWD1 and AIN76A diets. In the SI of control mice, tamoxifen treatment led to a robust expression of the dTomato reporter specifically in *Lyz*^+^ and *c-Kit*^+^ Paneth cells at the crypt base (Fig. 3C and Supplementary Fig. 3B - D), while dTomato was absent in *OLFM4*^+^ ISCs (Supplementary Fig. 3E). In the colon, dTomato was specifically expressed in *c-Kit*^+^ DCS cells, although the labelling efficiency was much lower when compared with the SI (Fig. 3C and Supplementary Fig.3 B and C).

For both Paneth and DCS cell lineages, the frequency at which lineage tracing was observed gradually and proportionally increased with the length (from 1 week to 1 month) of the NWD1 dietary treatment, while after 3 months, no further increase was observed (Fig. 3C and D). In both the small and large intestine, the lineage tracing ribbons emerged from *c-Kit*^+^ cells located at the base of the crypt, where PCs and DCS cells are localized (Supplementary Fig. S3F). Further immunostaining of the dTomato^+^ ribbons in NWD1-fed mice with antibodies specific for enterocytes (Villin), goblet (Muc2), tuft (Dclk1), and enteroendocrine (ChrgA) cells, confirmed that the mouse Western-style diet induces multipotency in *c-Kit*^+^ cells (Fig. 3E).

Collectively, these findings support that the WSD, similar to what previously observed with severe inflammation, rapidly suppresses *Lgr5*^+^ ISCs while simultaneously activating multipotent stem cell features in committed secretory lineages, namely PCs and DCS cells in the small and large intestine, respectively.

### NWD1 enhances colon tumor onset

It was previously reported that normal (i.e. not genetically modified) C57Bl/6 mice fed with NWD1 for longer intervals (18-24 months) develop benign and malignant neoplasms in the colon^25^. To confirm the NWD1-driven increased susceptibility to colonic tumors, *Cdx2*^CreERT2^; *Apc*^fl/+^ mice were fed for 1 months with the AIN76A and NWD1 diets before administering tamoxifen to induce monoallelic deletion of the adenomatous polyposis coli (*Apc*) tumor suppressor gene in *Cdx2*-expressing epithelial cells, i.e. in virtually all colonic lineages. Of note, in this model, tumor onset requires the acquisition of a second somatic hit at the wild-type allele, leading to loss of Apc function in actively dividing cells. Six months after tamoxifen-induced generation of the first somatic hit, only one mouse of the AIN76A-fed group developed a single hyperplastic lesion, while multiple polypoid and hyperplastic lesions all mice fed the NWD1 diet (Fig. 3F and Supplementary Fig. 3G). Notably, 9 months post induction mice maintained on NWD1 developed invasive colorectal adenocarcinomas, while we detected no malignant tumors in the AIN76A group (Fig. 3G and H).

These findings suggest that, despite the NWD1-mediated suppression of *Lgr5*^+^ ISCs, tumor formation is enhanced, most likely through the expansion or activation of alternative diet-induced stem cell populations, such as *c-Kit*⁺ deep crypt secretory (DCS) cells. Notably, early lesions of 6 months NWD1 fed mice contained Ki67⁺/c-Kit⁺ double-positive cells (Fig. 3I), indicating that DCS cells may contribute to the increased tumor burden observed in NWD1-fed animals.

### NWD1 causes persistent but reversible disruption of stem cell homeostasis

To determine whether suppression of *Lgr5*⁺ ISC lineage-tracing activity is permanent—persisting even after dietary reversal—or instead dependent on continued exposure to the Western-style diet, we first maintained mice on the two diets for an extended period (16 months). No lineage tracing from *Lgr5*⁺ ISCs was detected in NWD1-fed animals (Supplementary Fig. S3H), showing sustained suppression of resident stem cell activity for as long as the WSD was maintained. In contrast, Paneth cells retained lineage-tracing activity throughout the entire 16-month feeding period (Supplementary Fig. S3J), demonstrating that the diet-induced alterations of the ISC niche persist during chronic exposure to the rodent Western-style diet.

We then asked whether these perturbations of the intestinal stem cell niche can be reversed by switching mice from NWD1 back to the AIN76A control diet. As shown in Fig. 3J-L the lineage-tracing capacity of *Lgr5*⁺ ISCs was fully restored within 4 weeks of dietary reversal, whereas we did not observe any lineage tracing of *c-Kit*⁺ cells upon switching mice back to AIN76A diets.

### Western style diet induces inflammatory and genotoxic stress in *c-Kit*^+^ DCS cells

We next investigated diet-induced transcriptomic changes by single-nucleus RNA sequencing (snRNA-seq) of colonic tissues from mice fed either AIN76A or NWD1 diets. Following pre-processing, and sub-clustering of epithelial cells, we obtained transcriptomic profiles from a total of 39,400 epithelial cells representing all major colonic epithelial lineages (Supplementary Fig. S4A).

We first assessed whether NWD1 altered the cellular composition of the colonic epithelium. Although overall lineage distributions were largely preserved, mice fed for 1 month with NWD1 showed a trend toward increased proportions of secretory progenitor, enteroendocrine, and deep crypt secretory (DCS) cells (Supplementary Fig. S4B).

To unbiasedly evaluate which epithelial subpopulations are most sensitive to dietary stress, we applied the cell-type prioritization algorithm Augur^26^ to our single-cell RNA-sequencing dataset. Secretory progenitor and DCS cells emerged as one of the top cellular responders to NWD1 feeding (Supplementary Fig. S4C), indicating that DCS are among the cell types transcriptionally most responsive to dietary stress.

To examine the *c-Kit*⁺ DCS population, identified in our lineage-tracing experiments in greater detail, we subclustered *c-Kit*⁺ epithelial cells based on previously defined lineage-specific transcriptional signatures^24^ (Fig. 4A). Also here, the general distribution was maintained, although the NWD1 feeding resulted in minor changes in the proportion of *c-Kit*⁺ DCS cells which increased from 21.1 to 26.8% while *c-Kit*⁺ goblet cells decreased from 51.3 to 46.6% (Supplementary Fig. S4D). Cell-cycle analysis on *c-Kit*^+^ IECs demonstrated a shift toward increased proliferation within the *c-Kit*⁺ DCS compartment, with a greater proportion of cells entering S and G2/M phases in 1-month NWD1-fed mice compared with AIN76A-fed controls (Fig. 4B and Supplementary Fig. S4E). Of note, a similar increase in cycling cells was observed in a small *c-Kit*⁺ subpopulation (<2% of all *c-Kit*⁺ cells) expressing enteroendocrine cell markers (Fig. S4D and E).

**Figure 4:**
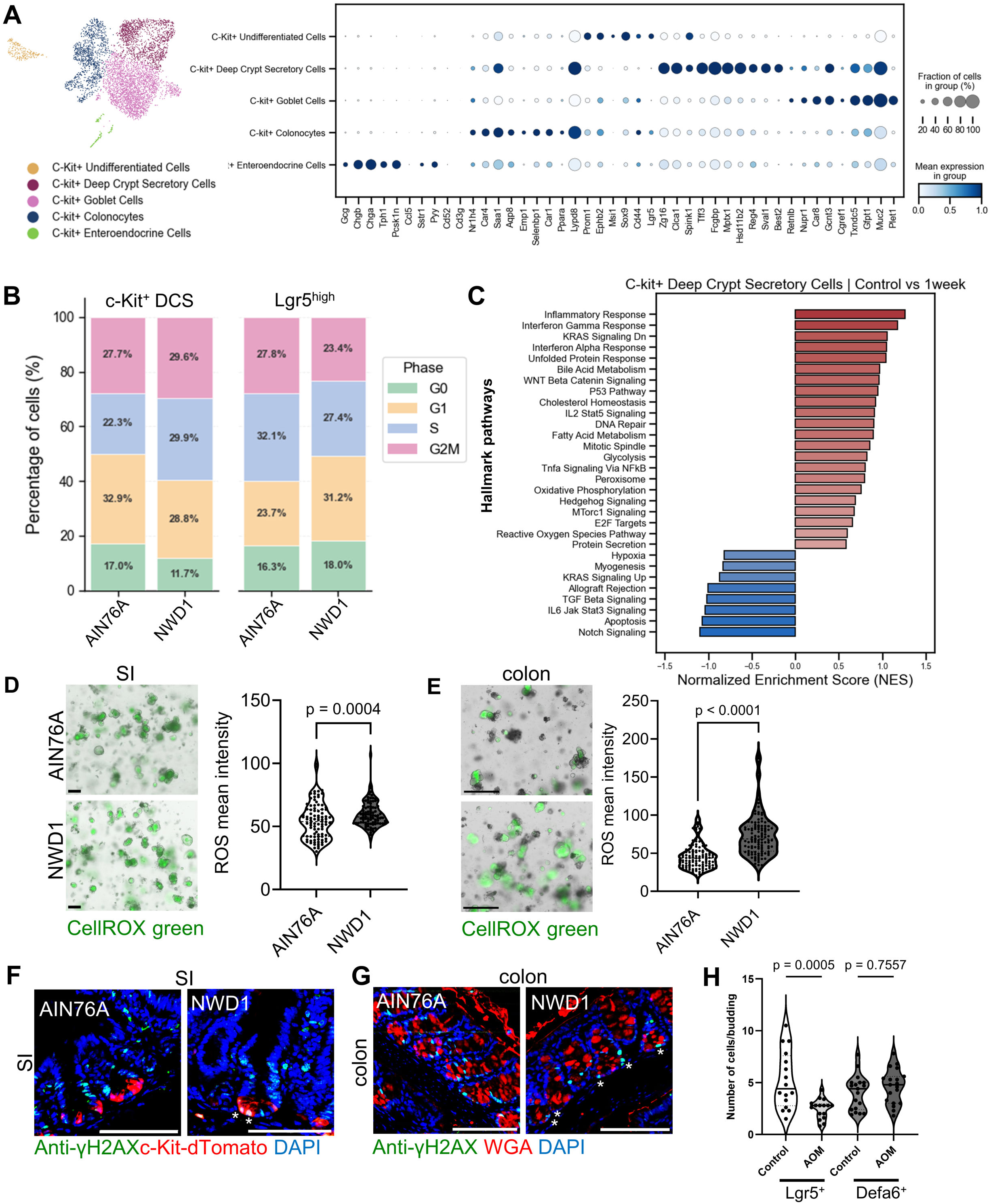
NWD1 induces inflammatory and genotoxic stress in *c-Kit*^+^ DCS cells. **A)** Uniform Manifold Approximation and Projection (UMAP) plot depicting 5 distinct clusters of the *c-Kit*^+^ cells (left) and their specific genetic profile (right). **B)** Bar plot of the distribution of *c-Kit*^+^ DCS cells and *Lgr5*^high^ ISCs of AIN76A and NWD1 fed mice, across cell cycle phases, scored using Seurat cell cycle gene signatures (Scanpy implementation). **C)** Bar plot summarizing the GSEA between *c-Kit*^+^ DCS cells of AIN76A and NWD1 fed mice. Pathways from the MSigDB Hallmark gene set collection are shown, filtered by |NES| > 0.5. No p-value cutoff was applied. **D** and **E)** Fluorescence microscopic ROS analysis of SI (D) and colonic (E) organoids derived from mice fed for 1 month with AIN76A or NWD1 using CellROX Green. Quantifications show mean ROS intensity per organoid and represents pooled data of 3 independent biological replicates (scale bar = 500µm for SI and 200 µm for colonic organoids). **F)** Confocal microscopy analysis of γ2HAX expressing dTomato labelled cells in the small intestine of 1-month AIN76A and NWD1 fed *c-Kit*^CreERT2^/*R26*^LSL-dTomato^ mice. Asterisks indicate γ2HAX^+^/dTomato^+^ cells (n = 3 mice per group, scale bars = 100 µm, nuclei were counterstained with DAPI). **G)** Confocal microscopy analysis of γ2HAX expressing WGA labelled cells in the colon of 1-month AIN76A and NWD1 fed *c-Kit*^CreERT2^/*R26*^LSL-dTomato^ mice. Asterisks indicate γ2HAX^+^/WGA^+^ cells at the base of colonic crypts (scale bars = 100 µm, nuclei were counterstained with DAPI). **H)** Quantification of GFP-labelled cells/crypt budding in *Lgr5*^EGFP-IRES-CreERT2^ and *Defa6*^Cre^/*Rosa*^mTmG^ organoids treated with AOM for 24 hrs (n= pooled data from 3 independent biological replicates). All data are mean ± s.d. and were analyzed by two-tailed Student’s *t*-test.

In contrast, clustering and analyzing *Lgr5*^high^ stem cells revealed a reduction in the proportion of cells in S and G2/M phases after 1 month of NWD1 feeding (Fig. 4B). Collectively, these snRNA-seq data are consistent with our lineage-tracing results and indicate that Western-style dietary exposure promotes proliferative expansion of *c-Kit*⁺ secretory populations, particularly DCS cells, while reducing proliferative activity within the *Lgr5*^high^ stem cell compartment.

To identify early molecular changes underlying this response, we performed Hallmark pathway analysis of *c-Kit*⁺ DCS cells isolated from mice fed NWD1 for 1 week. In addition to alterations in pathways regulating proliferation and cell fate, including activation of Wnt/β-catenin and mTORC1 signaling and suppression of TGF-β and Notch signaling, *c-Kit*⁺ DCS cells displayed enrichment of inflammatory response pathways together with increased expression of DNA repair and reactive oxygen species (ROS) response genes (Fig. 4C). These findings suggest that the newly proliferating DCS cells are exposed to both inflammatory and genotoxic stress. To validate these findings, we assessed oxidative stress and DNA damage in intestinal tissues from NWD1-fed mice. CellROX analysis revealed elevated ROS levels in both small intestinal and colonic organoids (Fig. 4D and E). Consistent with increased genotoxic stress, Paneth and DCS cells displayed nuclear staining for the DNA damage marker γH2AX (Fig. 4F and G), indicating activation of the DNA damage response in these secretory lineages. Together, these findings demonstrate that NWD1-induced intestinal inflammation is associated with enhanced proliferation and activation of stem cell-associated transcriptional programmes in *c-Kit*⁺ DCS cells, while simultaneously exposing these cells to oxidative and genotoxic stress. Despite this, pro-apoptotic genes were downregulated in *c-Kit*⁺ DCS cells from NWD1-fed mice, to the opposite of what was observed in Lgr5^high^ ISCs (Supplementary Fig. 4F). We then tested whether canonical *Lgr5*^+^ ISCs are more sensitive towards DNA-damage induced apoptosis then differentiated secretory cells, by exposing organoids of *Lgr5*^EGFP-IRES-CreERT2^ or *Defa6*^Cre^; *Rosa*^mTmG^ mice, labelling *Lgr5*^+^ ISCs or *Defa*6^+^ PCs, to the DNA damaging agent azoxymethane (AOM). While AOM treatment significantly reduced numbers of *Lgr5*^+^ ISCs, the number of *Defa6*^+^ cells remained unchanged (Fig. 4H), indicating their increased resistance to genotoxic stress. These results suggest that NWD1 promotes the expansion of proliferative *c-Kit*⁺ PC and DCS cell populations while increasing their resistance to DNA damage-induced apoptosis, potentially enabling the persistence of genetically stressed cells and thereby accelerating tumorigenesis.

Collectively, our findings demonstrate that NWD1 induces an inflammatory and genotoxic environment for intestinal epithelial cells, which suppresses proliferation and promotes apoptosis in *Lgr5*⁺ ISCs, while enhancing the stemness and survival of *c-Kit*⁺ DCS, even under conditions of DNA damage. This suggests that, in the context of a Western diet, alternative stem cell populations—rather than canonical ISCs—may drive CRC initiation, mirroring observations made during acute intestinal inflammation^12^.

### The enterotoxigenic *Bacteroides fragilis*, increases Wnt-signaling and drives intestinal stemness via *Bacteroides fragilis* toxin

Western diets have been shown to promote intestinal inflammation through alterations of the gut microbiome^27^. Consistent with this, 16S rRNA gene amplicon analysis of stool samples from NWD1-fed mice revealed marked shifts in microbiome composition, including significantly reduced microbial diversity, decreased abundance of several bacterial taxa (e.g. *Lactobacillaceae*, *Streptococcaceae*, *Lachnospiraceae*, and *Oscillospiraceae*), and enrichment of others (e.g. *Sutterellaceae* and *Tannerellaceae*) compared with AIN76A-fed controls (Fig. 5A and B and Supplementary Fig. 5A). These findings were further validated by semi-quantitative PCR analysis of stool DNA, which confirmed reduced levels of *Lactobacillus* spp. and increased abundance of *Sutterella* spp. in NWD1-fed mice (Fig. 5C).

**Figure 5:**
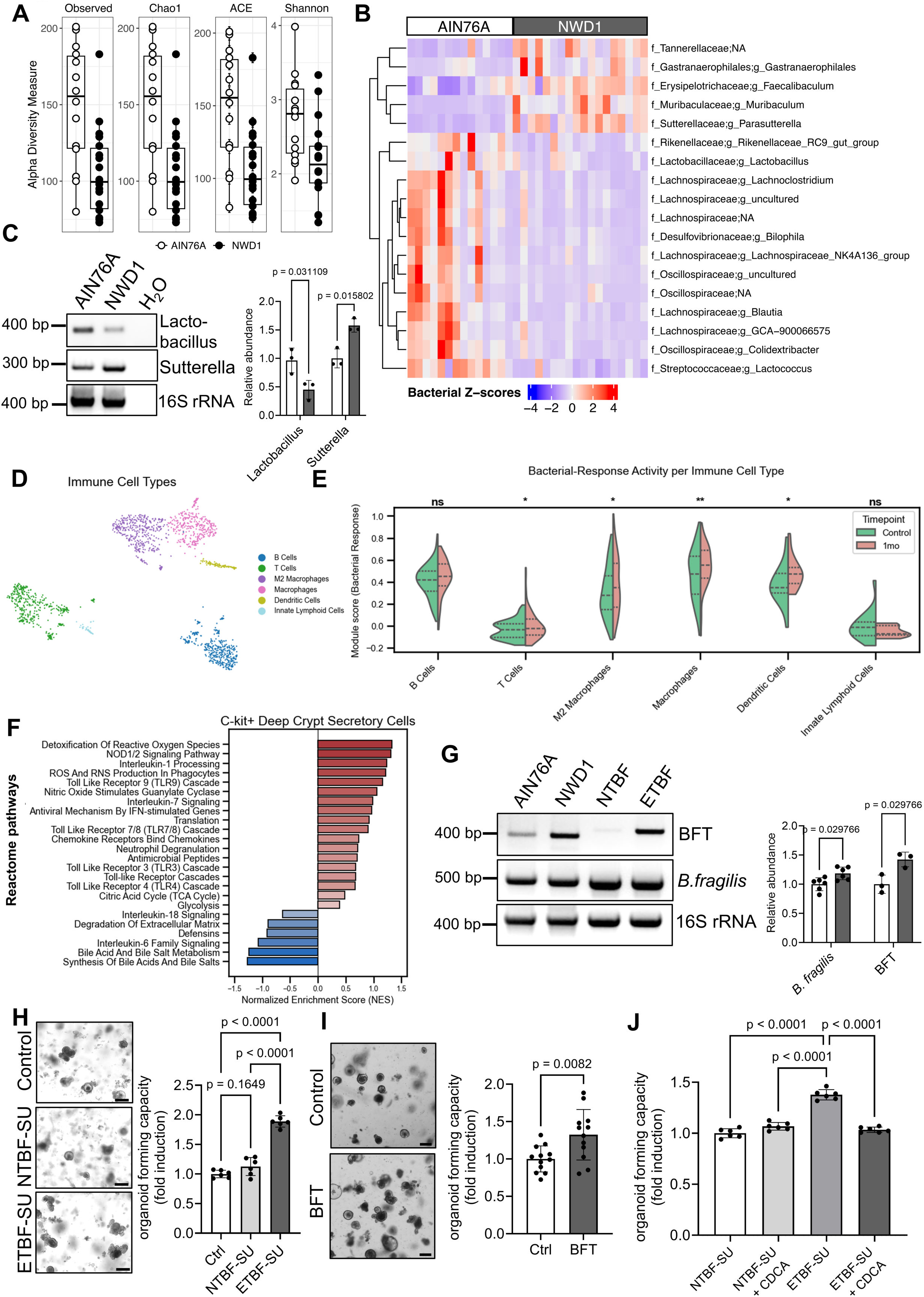
Enterotoxic *Bacteroides fragilis* is upregulated upon NWD1 and enhances intestinal stemness *ex vivo*. **A)** 16S rRNA gene amplicon sequencing of stool from different 3-months AIN76A and NWD1 fed mice showing reduced alpha-diversity in NWD1 fed mice using the indicated indices (n = 14 mice for AIN76A and 18 mice for NWD1). **B)** Z-score heatmap of differentially abundant microbes by diet detected by 16s RNA analysis in 3-month AIN76A and NWD1 fed mice (n = 14 for AIN76A and 18 for NWD1). **C)** Representative image of a PCR analysis of the indicated bacterial on stool-DNA from mice fed for 1 month with AIN76A and NWD1. Graphs show the fold change of intensity of *Lactobacillus* and *Sutterella* bands, normalized to the band intensities of the corresponding PCR bands of 16s primer set detecting all bacteria (n = 3 biological replicates). **D)** UMAP plot depicting distinct clusters of immune cells according to the transcriptional profile shown in Supplementary Fig. 4B (n= 3 mice per condition). **E)** Split violin plots showing a bacterial response module score across immune cell populations 1-month AIN76A and NWD1-fed mice. The module score was computed using Scanpy’s score_genes function on a curated set of bacterial response genes. Dashed lines indicate quartiles. Significance was assessed by two-tailed Welch’s t-test (n = 3 mice per condition). **F)** GSEA of *c-Kit*^+^ DCS cells from 1-month AIN76A and NWD1-fed mice using the Reactome 2022 pathway database, highlighting differentially enriched pathways (n = 3 individual mice per condition, pathways are filtered by |NES| > 0.5; no p-value cutoff applied. **G)** Representative image of a PCR analysis on stools from 1-month AIN76A and NWD1 fed mice using primers against *B. fragilis spp.* or specific to BFT. DNA isolated from non-enterotoxigenic (NTBF) or ETBF strains (DB_066 and DB05_075, respectively) were used as negative and positive control, respectively. Graphs show the quantification of *B. fragilis* and BFT band intensities normalized to the band intensities of the corresponding PCR bands of 16s primers detecting all bacteria (n = 3 biological replicates). **H)** Organoid multiplicities of SI crypts treated with organoid medium (control), or organoid medium supplemented with supernatants from NTBF or ETBF. (n = 3 biological replicates performed in duplicates, scale bar = 200µm). **I)** Organoid multiplicities murine SI crypts cultured in the absence or presence of recombinant *Bacteroides fragilis* toxin (BFT) (n = 4 biological replicates performed in triplicates, scale bar =200 µm). **J)** Organoid multiplicities of murine SI crypts cultured in organoid medium supplemented with NTBF or ETBF supernatants in the absence or presence of the BFT inhibitor CDCA (n = 4 biological replicates). All data are mean ± s.d. and were analyzed by two-tailed Student’s *t*-test (**C**,**G** and **I**) or one-way ANOVA (**H** and **J**) with Tuckey’s multiple comparison test.

We next explored our snRNA-seq dataset to determine whether these diet-induced microbiome alterations may be associated with the activation of the intestinal immune system in NWD1 fed mice. To this end, we subclustered and analyzed immune cell populations from AIN76A- and NWD1-fed mice (Fig. 5D and Supplementary Fig. S5B), revealing increased expression of genes associated with antibacterial immune responses (Fig. 5E), suggesting that WSD-induced inflammation is, at least in part, promoted by an altered microbiome.

In the intestinal epithelium both PCs and DCS cells, express Toll-like receptors (TLRs) and antimicrobial peptides and function as sensors of bacterial infection^28,29^. We therefore investigated whether DCS cells respond to WSD-induced microbiome alterations. Notably, our snRNA-seq analysis revealed increased activation of TLR-mediated signaling pathways, along with altered expression of antimicrobial peptides in DCS cells from NWD1-fed mice (Fig. 5F and Supplementary Fig. 5C), suggesting that WSD-associated microbial shifts are directly sensed by DCS cells.

We next examined *ex vivo* whether diet-induced alterations in the gut microbiota directly contribute to the observed effects on intestinal stem cell homeostasis. As lactate supports the oxidative metabolism and stemness of *Lgr5*^+^ ISCs^30^, we first addressed whether a reduction of lactate-producing microbes (i.e. *Lactobacillus* or *Lactococcus*) may underlie the suppression of canonical ISCs. Thus, we analyzed whether the *Lgr5*^+^ ISCs function in NWD1 fed mice can be restored *ex vivo* by supplementing the culture medium of freshly isolated crypts with lactate. However, we failed to observe any significant effect on the organoid forming capacity of both crypts from AIN76A- and NWD1-fed animals (Supplementary Fig. 5D).

Next, we analyzed the 16S rRNA gene amplicon sequencing data to identify bacterial taxa enriched in NWD1-fed mice that could potentially contribute to the diet-induced alterations of the ISC niche. While NWD1 feeding increased the abundance of the taxa *Muribaculum*, *Faecalibaculum*, and *Gastraanaerophilales* (Fig. 5B; Supplementary Fig. 5A), which have previously been associated with beneficial effects on gut health^31^, we also observed alterations in taxa with potential pathogenic properties. Notably, the relative abundance of *Sutterella* was significantly increased in NWD1-fed mice (Fig. 5B and C), and a trend toward increased *Bacteroides* abundance was detected, although this did not reach statistical significance (Supplementary Fig. 5E).

Among *Bacteroides fragilis* strains, enterotoxigenic *B. fragilis* (ETBF) has been strongly implicated in chronic intestinal inflammation and colorectal tumorigenesis. ETBF secretes the *B. fragilis* toxin (BFT), also known as fragilysin, a metalloprotease that binds Claudin-4 on epithelial cells and cleaves E-cadherin, thereby disrupting epithelial barrier integrity, impairing cell–cell adhesion, and promoting intestinal inflammation^32–35^. PCR analysis of stool samples from AIN76A and NWD1 fed mice confirmed an increase *B. fragilis*, particularly ETBF (Fig.5G). As BFT secreted from ETBF has been reported to activate β-Catenin signaling and to increase intestinal cell proliferation^36^, we hypothesized that BFT producing ETBF might exert direct effects on the ISC niche. Hence, we grew organoids from freshly isolated crypts in culture medium supplemented with supernatants of either ETBF bacterial cultures (ETBF-SU) or of non-enterotoxigenic *B. fragilis* cultures (NTBF-SU), which do not secrete the toxin. Only the supernatant of enterotoxigenic *B. fragilis* enhanced organoid growth (Fig. 5H) and increased β-Catenin activation and YAP-expression (Supplementary Fig. S5F). Of note, similar effects were observed with human colonic organoids cultured in medium supplemented with ETBF supernatants (Supplementary Fig. S5G and H). In contrast to ETBF, supplementing the culture medium of freshly isolated crypts with supernatants from *Sutterella wadsworthensis* cultures, failed to produce any significant effect on organoid formation capacity when compared to the culture medium lacking *S. wadsworthensis* supernatant (Supplementary Fig. S5I).

Next, we tested whether BFT itself has the capacity to increase organoid growth and cultured murine intestinal organoids with recombinant BFT, which improved organoid formation in a similar fashion to the ETBF supernatants (Fig. 5I). To validate these results, we supplemented the ETBF supernatants with chenodeoxycholic acid (CDCA), a direct BFT-inhibitor^37^, and observed the loss of the positive effects of ETBF-SU on organoid growth (Fig. 5J).

### NWD1-induced stem cell reprogramming is dependent on the microbiome and is mimicked by ETBF infection

Next, we analyzed whether ETBF, in addition to augmenting organoid formation, also alters ISC homeostasis. To this aim, we cultured crypts from *Lgr5*^EGFP-IRES-CreERT2^ mice in culture media supplemented either with NTBF- or ETBF-SU, or with recombinant BFT. As shown in Figure 6A, both ETBF-SU and the recombinant BFT had a significant detrimental effect on *Lgr5*-expressing cells within the organoids.

**Figure 6:**
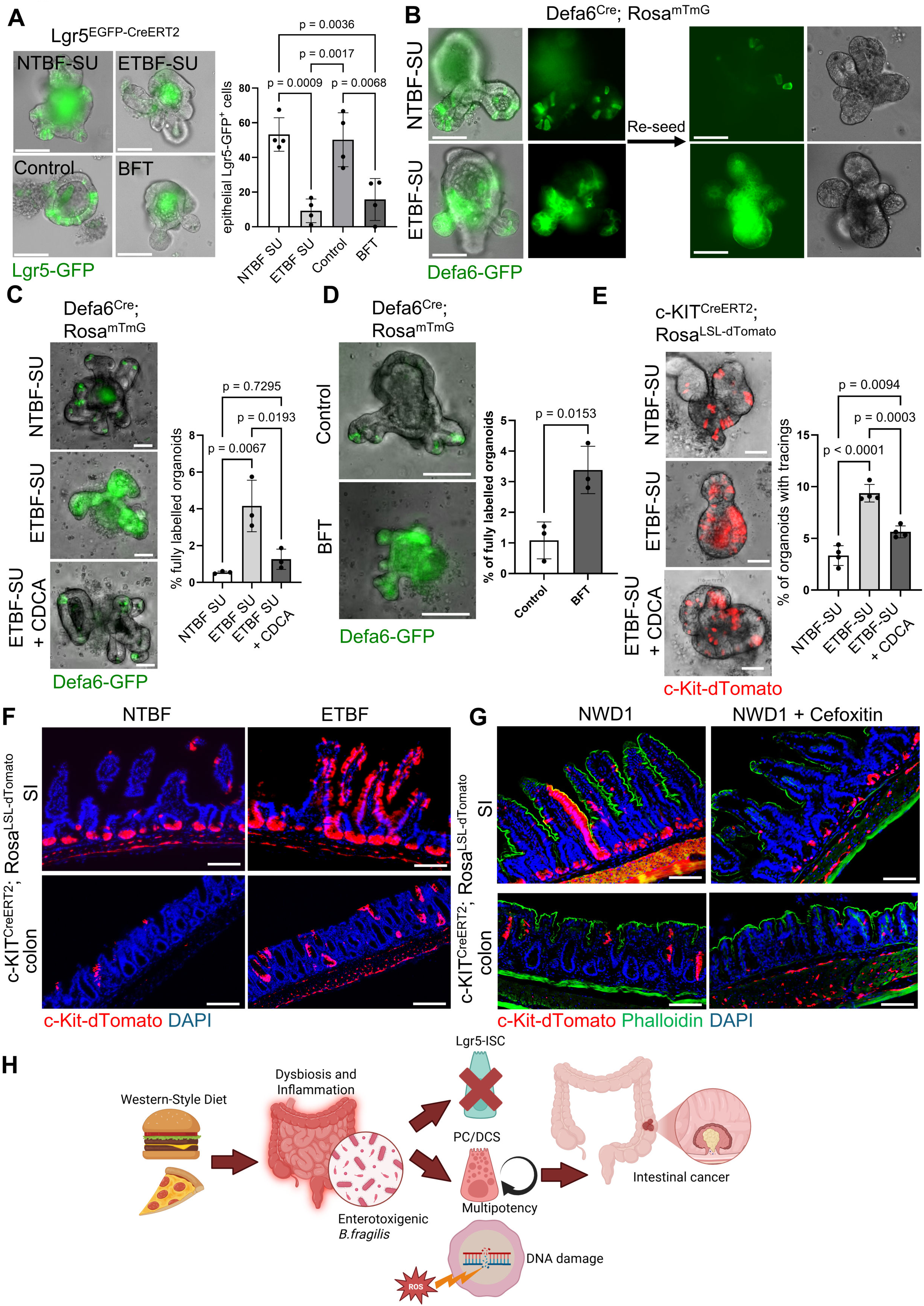
Western-style diet induced cellular plasticity is dependent on the gut microbiome and is mimicked by enterotoxigenic *B. fragilis in vivo*. **A)** Fluorescence microscopy analysis of SI organoids derived from *Lgr5*^EGFP-IRES-CreERT2^ mice treated with regular organoid medium or organoid medium supplemented with NTBF or ETBF supernatants or recombinant BFT. Graph shows the percentage of organoids with epithelial or GFP expression (n= 4 biological replicates, scale bars = 100 µm). **B)** Fluorescence microscopy analysis of SI organoids derived from *Defa6*^Cre^/*Rosa*^mTmG^ mice treated with organoid medium supplemented with NTBF or ETBF supernatants. Left: organoids before passaging, right: organoids after passaging. (n= 3 biological replicates, scale bars = 100 µm.) **C)** Fluorescence microscopy analysis of passaged SI organoids derived from *Defa6*^Cre^/*Rosa*^mTmG^ mice treated with organoid medium supplemented with NTBF or ETBF supernatants, in the absence or presence of CDCA. Graph shows the percentage of fully GFP labelled organoids (n= 3 biological replicates, scale bars = 50 µm). **D)** Fluorescence microscopy analysis of *Defa*6^Cre^/*Rosa*^mTmG^ SI organoids passaged after treatment with organoid medium or medium supplemented with recombinant BFT. Graph shows the percentage of organoids with epithelial or GFP expression (n= 3 biological replicates, scale bars = 50 µm). **E)** Fluorescence microscopy analysis of tamoxifen treated colonic organoids derived from *c-Kit*^CreERT2^/*Rosa*^dTomato^ mice cultured in organoid medium supplemented with NTBF or ETBF supernatants, in the absence or presence of CDCA. Graph shows the percentage of organoids with detectable lineage tracing (n= 4 biological replicates, scale bars = 50 µm). **F**) Confocal microscopy images of intestinal tissues of tamoxifen treated *c-KIT*^CreERT2^; *Rosa*^LSL-dTomato^ mice, infected with NTBF or ETBF (n = 3 mice per condition, scale bar = 100 µm, tissues were counterstained with phalloidin and DAPI). **G)** Confocal microscopy images of intestinal tissues of tamoxifen treated *c-KIT*^CreERT2^; *Rosa*^LSL-dTomato^ mice fed for 1 month with NWD1 diets in the presence or absence of Cefoxitin (n = 3 mice per condition, scale bar = 100 µm, tissues were counterstained with Phalloidin and DAPI). **H)** Schematic of Western-Style diet induced intestinal stem cell alterations. All data are mean ± s.d. and were analyzed by one-way ANOVA (**A**, **C** and **D**) with Bonferroni’s multiple comparison test or two-tailed Student’s *t*-test (**D**).

We then tested whether *B. fragilis* culture supernatants can induce the reprogramming of PCs and their acquisition of stem-like features in small intestinal organoids from *Defa6*^Cre^; *Rosa*^mTmG^ mice, where PCs and their potential daughter cells are marked by GFP^7,38^. We detected GFP-positive ribbons arising from the budding structures in organoids cultured with ETBF-SU, while GFP-positive cells were restricted to single PCs within the budding structures of organoids grown in NTBF-SU supplemented culture medium (Fig. 6B). Passaging the *Defa6*^Cre^; *Rosa*^mTmG^ organoids cultured in ETBF-SU medium, resulted in fully GFP-labelled organoids, an effect rarely observed when organoids were cultured in NTBF-SU medium (Fig. 6B-C), substantiating that ETBF-driven enhancement of PC multipotency *ex vivo.* Of note, we were able to inhibit the ETBF-driven induction of PC stemness in *Defa6*^Cre^; *Rosa*^mTmG^ organoids by supplementing the culture medium with CDCA (Fig. 6C). *Vice versa*, supplementation of the organoid culture medium with recombinant BFT resulted in lineage tracing from Paneth cells (Fig. 6D), showing that the toxin itself can induce PC multipotency.

To assess whether ETBF supernatants, analogous to the reprogramming of small intestinal PCs, can induce stemness of colonic DCS cells, we generated and tamoxifen-treated colonic organoids from *c-Kit*^CreERT2^; *Rosa*^LSL-dtomato^ mice, and cultured them with NTBF-SU or ETBF-SU media. We observed increased lineage tracing frequency in ETBF-SU organoids, which was again significantly reduced by CDCA (Fig. 6E), again pointing to the key role played by BFT in the reprogramming of both PCs and DCS cells.

To confirm the organoid results *in vivo*, tamoxifen-treated *c-Kit*^CreERT2^; *Rosa*^LSL-dtomato^ mice were infected with the NTBF and ETBF strains. ETBF treatment strongly enhanced the lineage tracing frequency of *c-Kit*^+^ cells in both the SI and colon when compared with NTBF-treated mice (Fig. 6F). This led us to hypothesize that the effects of WSD on stem cell homeostasis are mediated, at least in part, by microbiome alterations and the expansion of cancer-associated bacteria such as ETBF. If so, antibiotic treatment should attenuate these effects in NWD1-fed mice. Accordingly, treating *c-Kit*^CreERT2^; *Rosa*^LSL-dtomato^ mice with cefoxitin, a broad-spectrum antibiotic used to eradicate *B. fragilis*^39^ (Supplementary Fig. S6A), suppressed the capacity of NWD1 to induce lineage tracing from *c-Kit*^+^ cells, and to activate YAP and β-catenin signaling in small intestinal Paneth cells (Fig. 6G and Suppl. Fig. S6B and C). Last, cefoxitin reduced the number of Ly6G positive neutrophils and CD3^+^ T cells in the intestinal mucosa of NWD1-fed mice and suppressed the silencing of *Lgr5*^+^ ISCs (Supplementary Fig. S6D-F), suggesting that the WSD-mediated microbial changes are likely upstream of intestinal inflammation and stem cell reprogramming.

In summary we show that WSD rapidly and reversibly reshapes intestinal stem cell homeostasis by suppressing canonical *Lgr5*⁺ ISCs while inducing stem-like properties in differentiated secretory cells. Despite reduced *Lgr5*⁺ ISCs activity, WSD enhances colorectal tumor initiation, suggesting that facultative stem cell populations can act as alternative cells of origin. ISC reprogramming is associated with chronic low-grade inflammation and microbiome alterations, including expansion of enterotoxigenic *Bacteroides fragilis*, whose toxin autonomously can promote epithelial plasticity and stemness. Together, our findings establish a functional link between diet, microbiota-driven inflammation, stem cell reprogramming, and colorectal cancer initiation (Figure 6H).

## Discussion

In homeostasis, the intestinal epithelium is sustained by rapidly cycling *Lgr5*⁺ ISCs. Yet it exhibits remarkable plasticity through recruitment of facultative stem cell populations upon inflammation and injury^3,40^. While such regenerative programs have been mainly characterized in acute tissue damage, whether chronic environmental exposures engage similar mechanisms is yet unclear. Here, we show that a rodent diet mimicking the composition of Western-style diet in man is pro-inflammatory, underlies specific microbiome alterations, and ultimately results in the rapid and reversible reprogramming of the intestinal stem cell niche by suppressing canonical *Lgr5*⁺ ISCs while activating alternative stem cell states derived from secretory lineages.

A central insight of our study is the uncoupling of canonical ISC activity from overall epithelial stemness and tumorigenesis in the context of Western dietary habits.

Despite early and sustained inhibition of *Lgr5*⁺ ISC proliferation, WSD paradoxically augments Wnt and YAP signaling, organoid-forming capacity, and tumor initiation. This demonstrates that tissue stemness is not only maintained but functionally amplified through alternative cellular sources. Specifically, we show that small intestinal PCs and their colonic counterparts, DCS cells, re-enter the cell cycle and acquire multipotency to drive a regenerative response against diet-induced inflammatory insults, reminiscent of their function in the context of acute inflammation or injury^5,6,8^.

However, we believe that the trade-off of this adaptive response is a reason why Western-style dietary habits represents the main etiologic risk factor for colon cancer. The novel stem-like cells originating from *c-Kit*⁺ secretory lineages, among other committed cell types, exhibit increased proliferation and, in the presence of elevated reactive oxygen species, undergo DNA damage and yet downregulate pro-apoptotic pathways. The latter will not only favor the accumulation of somatic mutations, but will also activate DNA damage response and epigenetic reprogramming, consistent with models where regeneration-associated plasticity increases susceptibility to tumor initiation^9–12,41^. Our findings therefore support a model where pro-inflammatory nutrients expand the pool of stem-like cell targets for mutations, thus providing a mechanistic link between diet and CRC risk. Accordingly, NWD1 increases the relative frequency of rate-limiting and tumor-initiating events, notwithstanding the suppression of canonical *Lgr5*^+^ ISCs, similar to what we observed in inflammation-associated CRCs^12^. In this scenario, inflammation and dysbiosis play a central role in mediating the diet-induced epithelial reprogramming of the intestinal stem cell niche. WSD results in profound microbial shifts, characterized by reduced microbial diversity and expansion of opportunistic microbes, alongside the activation of antibacterial immune responses. Eradication of the microbiome by means of antibiotics largely abrogates both inflammation and stem cell reprogramming, positioning the gut microbiome upstream of epithelial changes. These observations are consistent with prior findings demonstrating that microbial signals regulate ISC function and niche composition^42,43^, and further support a model where diet reshapes epithelial homeostasis through inflammation and microbiome modification.

Within this altered microbial landscape, we identified the enterotoxigenic *Bacteroides fragilis* (ETBF) as a key effector linking gut microbes to stem cell plasticity. ETBF and its toxin BFT, suppress *Lgr5*⁺ ISCs while promoting proliferation and multipotency of secretory PC and DCS cell lineages in the small and large intestine, respectively, as demonstrated here both *ex vivo* and *in vivo*. The ETBF-mediated activation of the Wnt/β-catenin pathway is consistent with its established role in modulating epithelial signaling and promoting tumorigenesis: ETBF contributes to the reprogramming of the *c-Kit*^+^ secretory lineages by directly activating stem cell-related pathways in *c-Kit*^+^ Paneth and DCS cells^32,36,44–46^. Our findings suggest that ETBF contributes to colorectal cancer risk not only by promoting chronic inflammation but also by directly regulating stem cell identity and epithelial plasticity. The ETBF-derived toxin BFT is known to disrupt epithelial barrier integrity through cleavage of the E-cadherin ectodomain, thereby triggering Wnt/β-catenin signaling^35,36,47^. By actively triggering stemness pathways in lineage-committed, mutation-prone PCs and DCS cells, ETBF expands the pool of tumor-competent target cells. This unique mechanism explains how the bacterium promotes colorectal carcinogenesis without directly inducing a distinct bacterial mutational signature^48^.

The reversibility of WSD-induced phenotypes provides an important conceptual insight. Restoration of a normal diet rapidly rescues *Lgr5*⁺ ISC activity and suppresses *c-Kit*⁺-driven stemness, indicating that epithelial reprogramming is dynamically maintained and dependent on continuous environmental input. This aligns with emerging evidence according to which ISC identity is highly plastic and responsive to niche-derived signals^40^, and suggests that dietary and microbiome-targeted interventions may have therapeutic potential even after prolonged exposure.

We acknowledge that there are limitations to our study. While ETBF emerged as a key mediator, WSD-induced microbiota alterations are complex, and additional microbial species and metabolites may contribute to the observed phenotype. Similarly, we cannot exclude that additional cell types, previously reported to acquire multipotency upon acute tissue injury, may regain stem cell capacities upon exposure to WSD, similar to *c-Kit*^+^ Paneth and DCS cells, thus further increasing the pool of cell targets for tumor initiation.

In summary, we identified Western-style diet as a potent regulator of intestinal stem cell identity, acting through inflammation- and microbiome-dependent mechanisms to promote the reprogramming and expansion of alternative stem cell populations. By linking WSD-induced microbial changes to epithelial plasticity, genotoxic stress, and stem cell reprogramming, and by identifying ETBF as a targetable mediator, our findings provide a conceptual framework connecting diet, microbiota, and CRC risk. These insights highlight the intestinal stem cell niche as a dynamic interface integrating environmental and microbial signals, with important implications for cancer prevention and intervention.

## Methods

### Mice

The following inducible Cre strains were used: *Lgr5*^CreERT2-EGFP^ (Jackson Laboratories, cat. no. 008875)^3^, *c-Kit*^CreERT2^ (kindly provided by Dieter Saur)^17^, p*Lys*^CreERT2^ (kindly provided by Hans Clevers)^49^, *Defa6*^Cre^ (kindly provided by Richard S Blumberg)^38^. For lineage-tracing experiments, mice were crossed with *R26*^LSL-YFP^ mice (Jackson Laboratories, cat. no. 006148), *R26*^LSL-tdTomato^ mice (Jackson Laboratories, cat. no. 007908) or *R26*^mT/mG^ mice (Jackson Laboratories, cat. no. 007676). *CDX2*^CreERT2^; *Apc*^Fl/Fl^ mice were purchased from (Jackson Laboratories, Strain #:035169, Wildtype for Kras<tm4Tyj>, Homozygous for Apc<tm1Tno>, Homozygous for Tg(CDX2-cre/ERT2)752Erf). All mice used here were inbred C57BL6/J and bred and maintained in the University of Marburg Mouse Facility or in the Erasmus MC animal facility (EDC) under conventional specific pathogen-free conditions in individually ventilated cages with a 12-hour/12-hour light–dark cycle. For all experiments, mice were randomly assigned to experimental groups after matching for sex, age of 4–6 weeks and genotype. All protocols involving animals were approved by the German Animal Welfare Act and in accordance with the local animal welfare authority and committee (Regierungspräsidium Giessen) or the Dutch Animal Experimental Committee and in accordance with the Code of Practice for Animal Experiments in Cancer Research established by the Netherlands Inspectorate for Health Protections, Commodities and Veterinary Public Health.

For diet experiments, mice were fed as indicated in the individual experiments, with AIN-76A or NWD1 diets^50^ (Altromin, formulations are based on diets D10011 (AIN-76A Rodent Diet, Research Diets) and D16378C (NWD1) Newmark Stress Diet C, Research Diets) and are provided in the Supplementary Data File). In brief, NWD1 is a rodent diet based on AIN-76A control diet, designed to mimic the suggested colon cancer risk factors in humans. Increased dietary fat (40% of total calories or 20% of weight of diet); lower dietary calcium (0.5 mg/g), reduced vitamin D_3_ (0.11 IU/g diet, approximately equivalent to 50 IU/day in a human 2000 kcal diet). The levels of folic acid, were similarly reduced to the inadequate levels of some human diets, approximately to the lower one-fourth of the average human diets in the USA. Details for these evaluations and basis for the diet concentrations used have been reported previously^16^. To analyze the effect of experimental diets on CRC onset CDX2CreERT2; ApcFl/Wt mice were fed for 1 month with AIN76A or NWD1 diet and injected intraperitoneal with tamoxifen (1 mg dissolved in 100% ethanol and subsequently in corn oil; Sigma, cat. nos. T5648 and C8267). The mice were then fed for up to additional 9 months with the respective diets and then analyzed for intestinal tumor formation. To ablate intestinal microbiota, mice received cefoxitin, administered in autoclaved drinking water at a concentration of 0.5mg/ml. Antibiotic-containing water was replaced every 48 hours to ensure sustained antibiotic activity. The treatment was maintained in parallel with the feeding of NWD1 for 4 weeks. To inoculate mice with NTBF and ETBF, mice were gavaged once with the non-enterotoxigenic *B. fragilis* strain DSM 2151 (DSMZ, cat. No. 2151, Type strain) or the enterotoxigenic *B. Fragilis* strain TT12^51^ (kindly provided by Victor Sourjik, positivity for *B. fragilis* toxin was verified by sequence blast (blast report is provided in the Supplementary Data File)) at a density of 1×10^9^CFU/mL resuspended in drinking water, independent of weight.

### *In vivo* lineage tracing experiments

To induce C-recombinase mice were injected intraperitoneal with tamoxifen (1 mg dissolved in 100% ethanol and subsequently in corn oil; Sigma, cat. nos. T5648 and C8267) once or three times on 3 consecutive days. 7-10 days after the last tamoxifen injection, intestinal tissues were harvested for lineage-tracing analysis. Tissue samples were first dissected and washed with PBS buffer, and then fixed for 1h at 25°C with 4% PFA solution, followed by sequential cryoprotection in 15% sucrose for 3 hours at 4°C and 30% sucrose overnight at 4°C. Tissues were individually embedded in O.C.T. (Optimal cutting temperature compound; Science Services, SA62550-01) and stored at −80°C. Cryosections were cut at 10-12μm thickness using a Leica CM3050S cryostat. Cryosections were thawed at room temperature for 10 min and washed three times with PBS containing 0.2% Ecosurf™ SA-9 (PBST, Carl Roth, 09822.3). For Lgr5-GFP-dTomato reporter mice, sections were counterstained with 4,6-diamidino2-phenylindole (DAPI; Sigma, D9542-1MG) only, for 30 min at room temperature. For c-Kit lineage tracing experiments, sections were stained with DAPI and Alexa Fluor 488-conjugated phalloidin (1:200, Invitrogen, A12379), for 30 min at room temperature. After three additional 5min PBS washes, sections were mounted using Dako Fluorescence Mounting medium (Agilent Technologies Inc., S3023802). Images were acquired using a Zeiss LSM 700 laser scanning confocal microscope. Images were processed with ImageJ. A lineage trace was considered positive when ≥5 contiguous dTomato-labeled cells were observed extending from the crypt into the villus. For Lgr5-GFP-dTomato mice, GFP-positive crypts were categorized as either lacking dTomato labeling (GFP⁺ dTomato⁻), meaning no lineage tracing, or showing dTomato positive lineage tracing (GFP⁺ dTomato⁺). The percentage of lineage tracing was calculated as:

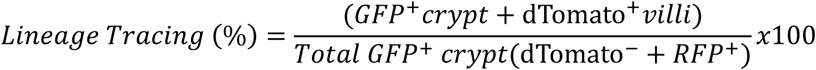

For c-Kit and pLys lineage tracing, dTomato-positive or YFP-positive crypts, respectively, were identified and assessed for lineage tracing into the villus compartment. dTomato⁺ crypts were categorized as either showing villus tracing or lacking tracing. The percentage of lineage tracing was calculated as:

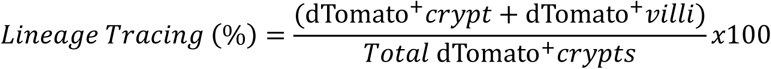

To ensure representative sampling, 100-250 crypts per section were quantified. Three different sections per intestinal segment (duodenum, jejunum, ileum, and colon) were analyzed per animal. A minimum of three animals per experimental condition and timepoint were evaluated.

### Immunohistochemistry

Tissues were fixed in 4% PFA overnight at 4°C and embedded in paraffin. The 4-8µm sections were deparaffinized by sequential immersion in xylene (2 x 5mins), followed by graded ethanol series (100%, 96%, 80%, 70%; 5mins each), and rehydration in deionized water for 10mins. Antigen retrieval was performed in a pressure cooker at pH 9 with either Tris-EDTA buffer or citrate buffer (1x), for 10 min. After cooling to room temperature, sections were rinsed in deionized water and endogenous peroxidase activity was quenched by incubation with 3% H₂O₂ (Merk, 1072091000) for 10mins. Sections were blocked with 4% goat serum (GS, Thermo Fisher Scientific, 16210064) in PBST for 1.5 hours at room temperature, then incubated overnight at 4°C with the following primary antibodies: β-catenin (Cell Signaling Technology, 8814); YAP (Cell Signaling Technology, 14074T); and Ki67 (BioLegend, 652402). Sections were washed three times in 2% GS in PBST, then incubated for 1.5 hours at room temperature with the following biotinylated secondary antibodies diluted in 2% GS in PBST: Anti-Rabbit (Vector Laboratories, BA-1000-1.5), and Anti-Rat (Vector Laboratories, BA-1000-1.5). Following three washes in 1% GS in PBST, sections were incubated with avidin-biotin complex (ABC) solution (Vector Laboratories, PK-6100) for 1 hour at room temperature in the dark. DAB chromogen solution (Vector Laboratories, SK-4100) was freshly prepared and applied to sections. Sections were counterstained with hematoxylin (Carl Roth, T865.2) for 30s and washed under running water for 5mins. Slides were dehydrated through graded ethanol (70%, 80%, 96%, 100%; 2mins each), cleared in xylene (2 x 5mins) and mounted with Pertex (Histolab, cat. no. 00811). Images were acquired using Leica LAS-X software.

### Immunofluorescence

For cryosections, tissues were first dissected from the mice and washed with PBS buffer, and then fixed for 1h at 25°C with 4% PFA solution, followed by sequential cryoprotection in 15% sucrose for 3 hours at 4°C and 30% sucrose overnight at 4°C. Tissues were individually embedded in O.C.T. (Optimal cutting temperature compound; Science Services, SA62550-01) and stored at −80°C. Cryosections were cut at 10-12μm thickness using a Leica CM3050S cryostat. For FFPE sections, tissues were harvested and washed with PBS buffer, and then fixed in 4% PFA overnight at 4°C. Tissues were subsequently embedded in paraffin using a Medite Tissue Embedding System (**#**TES99). Once solidified, paraffin blocks were sectioned at 4μm thickness using a Leica HistoCore MULTICUT microtome. Immunofluorescence was performed using antibodies against: Ki67 (1:200, Biolegend, 652402), c-Kit (1:100, Cell Signaling Technologies, 3074S), γH2A.X (1:200, Cell Signaling Technology, 9718T), CD3 (1:200, Abcam, AB16669), CD11C (1:200, Cell Signaling Technologies, 97585S), Ly6G (1:100, Cell Signaling Technology, 87048T), TNFα (1:200, Cell Signaling Technologies, 11948T), Lyz1 (1:500, Santa Cruz, sc-518012), OLFM4 (1:200; Cell Signaling Technologies, 39141), Villin (1:200, Santa Cruz, sc58897), Muc2 (1:200, Abcam, ab272692), DCLK1 (1:200, Abcam, ab31704), ChrgA (1:200, Abcam, ab15160), and YAP (1:200, Cell Signaling Technologies, 14074T). Tissue sections were washed and blocked with 5% serum (goat - Thermo Fisher Scientific, 16210064; milk - Carl Roth, T145.3; or BSA - Carl Roth, T844.4) for 1-2 hours at room temperature, then incubated with the primary antibody overnight at 4 °C. Slides were washed twice with PBST and incubated for 2 h with the following secondary antibodies: Goat anti-Rabbit Alexa Fluor™ Plus 488 (1:200, Invitrogen, A-11008), Goat anti-Rabbit Alexa Fluor™ Plus 555 (1:200, Invitrogen, A-21429), Goat anti-Rat Alexa Fluor™ Plus 488 (1:200, Invitrogen, A-11006), Goat anti-Rat Alexa Fluor™ Plus 555 (1:200, Invitrogen, A-21434), Goat anti-Mouse Alexa Fluor™ Plus 488 (1:200, Invitrogen, A-11001), and WGA 555 conjugate (1:500; ATT Bioquest/ #25550). Tissues were counterstained with DAPI (Sigma Aldrich, D9542-1MG) and mounted in Dako Fluorescence Mounting Medium (Agilent Technologies Inc, S3023802) and imaged with a Zeiss LSM700 confocal microscope.

### Quantification of immunostaining

In Fig. 1E, the total number of Ki67 positive cells, per small intestinal crypt was analyzed and averaged. The percentage of Ki67^+^C-kit^+^ double-positive cells (Fig. 2B) was quantified by dividing the number of c-Kit/Ki67 double-positive cells by the total amount of c-Kit cells in the lower half of the colonic crypt. Only clearly detectable double-positive cells, as determined by the analysis of *z*-stacks, were counted. For each quantification, at least 30 crypts of at least 3 sections/mouse of 3 experimental mice per group were used. The number of CD3, Ly6G or TNF positive cells/villus was quantified by counting the total number of positive cells per villus on at least 30 villi distributed over 3 sections of 3 experimental mice per group. As in Supplementary Fig. S3F, CD11c positive individual cells could not be separated, we quantified the intensity of the CD11c immunostaining/villus of at least 100 villi distributed over 3 sections of 3 experimental mice per group. Individual villi were manually outlined using the freehand selection tool, and mean fluorescence intensity was measured within each selected region. All images were acquired using identical exposure settings, and fluorescence intensity measurements were performed using consistent threshold parameters across all experimental conditions.

### FACS

As for organoid reconstitution assays (ORA), isolated crypts were first digested to single cells after incubation with TrypLE (Life Technologies) and 2000 U/ml-1 Dnase (Sigma) for 10 min. at 37°C. Dissociated cells were passed through a 40 μm strainer, stained with an APC-conjugated CD24 antibody (clone M1/69 BioLegend), and, to exclude non-epithelial cells, with the following Ab’s for 30 min. at 4°C: BV421-conjugated TER119 (Biolegend), CD31 (Biolegend), CD45 (Biolegend). For PC analysis, an additional staining with a PE-conjugated CD117 antibody (Biolegend) was also performed. Cells were then sorted by FACS (BD FACSAriaIII; BD Bioscience). Single viable epithelial cells were gated by CD24 and side scatter (SSC), and by negative selection against the above BV421-conjugated Ab’s and with DAPI to discriminate live from dead cells. Detailed FACS gating strategy for the small intestine was performed as previously described^4^. Reconstitution of Lgr5-EGFPhi stem cells (purity > 99%) with PCs (CD24hiSSChi; purity > 98%) was performed by pelleting sorted cells at 300 g. for 5 minutes in Eppendorf LoBind Tubes, and by co-incubating them for 15 min. at 25°C as previously described^21^.

### Bacterial culture and supernatant generation

Supernatants of non-enterotoxigenic *B. fragilis* strain DB_066 and enterotoxigenic *B. fragilis* strain DB05_075 were kindly provided by Jens Puschhof, DKFZ, Heidelberg, Germany. For generation of *B. fragilis* supernatants, microbial strains were cultured in modified Gifu Anaerobic Media (mGAM) in an anaerobic chamber. For generation of *Sutterella* supernatants, *Sutterella wadsworthsensis* (DSM14016) was cultured as recommended by the provider. Supernatants were extracted after overnight culture and subsequently sterile filtered (0.22µm filter) for organoid culture use. Negativity or positivity for *Bacteroides fragilis* toxin (BFT) was verified via PCR using BFT specific primers. The non-enteroxigenic *B. fragilis* strain DSM 2151 (DSMZ, cat. No. 2151, Type strain) and the enterotoxigenic *B. Fragilis* strain TT12^51^ used for the *in vivo* experiments were kindly provided by Victor Sourjik, MPI, Marburg, Germany. Positivity of the *B. fragilis* strain TT12 for *B. fragilis* toxin was verified by sequence blast (blast report is provided in the Supplementary Data File). Bacteria were cultured anaerobically at 37 °C on pre-reduced Columbia blood agar and in supplemented brain heart infusion medium (yeast extract, vitamin K, hemin). Single colonies were expanded in liquid culture to late exponential phase (OD600 0.35–0.4), pooled, pelleted (2000 × g, 5 min), and washed in sterile PBS. Bacterial loads were determined by serial dilution and CFU enumeration following 48 h anaerobic incubation. Pellets were snap-frozen at −80 °C and resuspended in sterile water at 1 × 10⁹ CFU ml⁻¹ immediately prior to use. Mice received 100 µl by oral gavage (1 × 10⁸ CFU).

### Isolation of murine intestinal crypt and organoid culture

Intestinal organoids were generated as previously described^52^. In brief, mouse small and large intestines were flushed with cold PBS, removed from fat and scraped with a glass slide to remove intestinal villi. Resected sample were then sectioned into 5–10-mm segments and washed with cold PBS before incubation for 45 min at 4 °C in cold PBS supplemented with 2.5 mM EDTA. After an additional PBS wash, crypts were detached from the muscle layer by four to five rounds of harsh shaking with cold PBS and supernatant was collected into new separated falcons. The optimal fractions containing the highest crypt density and minimal debris were pooled and filtered through a 70μm cell strainer, washed with PBS and centrifuged 80-120g for 2min at 4°C, four to five times. Single crypts were manually counted. Crypts were seeded at a 1000 crypt/drop density, in 20μL drops of Matrigel (Corning), using pre-warmed 12-well plates. Plates were immediately transferred to a 37°C, 5% CO₂ incubator for 10min to allow Matrigel polymerization. Following solidification, 1 mL of organoid culture medium was gently added to each well.

### Organoid experiments

To assess the effect of experimental diets on organoid growth, mice were fed for the indicated times with AIN76A or NWD1 diet and crypts were isolated and seeded as described above. Organoid numbers and sizes were assessed on days 4-7 post-seeding on images acquired with Nikon Eclipse Ts2 fluorescence microscope. To analyze the effect of AIN76A and NWD1 on organoid formation of isolated *Lgr5*^+^ and *c-Kit*^+^ Paneth cells, isolated crypts were first digested to single cells after incubation with TrypLE (Life Technologies) and 2000 U/ml-1 Dnase (Sigma) for 10 min. at 37°C. Dissociated cells were passed through a 40 μm strainer, stained with an APC-conjugated CD24 antibody (clone M1/69, Biolegend, cat.no: 101814; RRID: AB_439716), and, to exclude non-epithelial cells, with the following Ab’s for 30 min. at 4°C: BV421-conjugated TER119 (Biolegend, cat.no: 563998), CD31 (Biolegend, cat.no: 563356), CD45 (Biolegend, cat.no: 563890; RRID: AB_2651151). For PC analysis, an additional staining with a PE-conjugated CD117 antibody (Biolegend, cat.no: 105808; RRID: AB_313217) was also performed. Cells were then sorted by FACS (BD FACSAriaIII; BD Bioscience). Single viable epithelial cells were gated by CD24 and side scatter (SSC), and by negative selection against the above BV421-conjugated Ab’s and with DAPI to discriminate live from dead cells. Detailed FACS gating strategy for the small intestine was performed as previously described^53^. Isolated *Lgr5*-EGFPhi stem cells (purity > 99%) or *c-Kit*^+^ Paneth cells (CD24hiSSChi; purity > 98%) were pelleted at 300 g. for 5 minutes in Eppendorf LoBind Tubes, resuspended and 2000 FACS isolated *Lgr5*^+^ ISCs or *c-Kit*^+^ Paneth cells were seeded in individual Matrigel drops and organoid forming capacity was analyzed on day 5-10 after seeding. To analyze the effect of AOM on *Lgr5*^+^ ISC or *Defa6*^+^ PC numbers, small intestinal organoids were split one day before treatment. The following day, the organoid culture medium was supplemented with 100 ng/mL azoxymethane (AOM). Individual organoids were tracked longitudinally by imaging the same organoids at baseline (day 0) and on days 1, 2, and 3 of AOM treatment using a Leica THUNDER Imager. Z-stacks were analyzed in LAS X Office, and *Lgr5*-EGFP⁺ and *Defa6*-GFP⁺ cells were manually counted at each time point. For measuring the effects of bacterial supernatants, metabolites and toxins on murine organoid growth, intestinal crypts were isolated as described above and cultured in organoid medium supplemented with lactate (20 mM, Sigma-Aldrich, cat. No.: 71729-5G), the recombinant *Bacteroides fragilis toxin* fragilysin (BFT) (0.1mg/mL, Sigma-Aldrich, cat. No.: C9377-100MG) or supernatant of bacterial cultures (1:100 dilution). Where indicated organoid cultures containing *B. fragilis* supernatants were supplemented with chenodeoxycholic acid (CDCA, 50µM, Cusabio, cat.no.: CSB-EP346537BDP.20). Organoid numbers and size were assessed on day 4-7 post-seeding. For examining the effects of *B. fragilis* supernatants on human colonic organoids (derived from healthy human colon and kindly provided by Jens Puschhof, DKFZ, Heidelberg, Germany), the organoids were disassociated into single cells and 30 000 cells were seeded and grown in human colon expansion medium as previously described ^54^ for 5 days until small organoids were formed. Subsequently, the organoids were exposed in triplicates to supernatant of ETBF and NTBF diluted 1:100 in differentiation media (ENR) ^54^ for another 6 days and then analyzed. As a negative control the bacterial growth medium (mGAM) was used 1:100 in ENR. For the *ex vivo* analysis of the effect of BFT, ETBF and NTBF on *Lgr5*^+^ ISC numbers, we isolated crypts from *Lgr5*^CreERT2-EGFP^ mice as described above. On day 3, organoid medium was supplemented with BFT (0.1mg/mL), or ETBF and NTBF supernatants (1:100 dilution) and after additional 2 days in culture organoids were analyzed and the number of GFP positive cells in the epithelial lining quantified. For the *ex vivo* lineage tracing analysis of Paneth cells, we isolated small intestinal crypts from *Defa6*^Cre^; *Rosa26*^mTmG^ as described above and where indicated we supplemented the organoid medium with BFT (0.1mg/mL), 1:100 dilutions of NTBF supernatants, ETBF supernatants or ETBF supernatants containing CDCA (50µM). Lineage tracing events were assessed on day 5 from culture and subsequently organoids were, passaged. 3-5 after passaging the percentage of fully traced organoids on the total number of GFP-labelled organoids was quantified. For the *ex vivo* lineage tracing analysis of *c-Kit*^+^ DCS cells, we isolated crypts from *c-Kit*^CreERT2^; *Rosa*^LSL-dTomato^ as described above. On day 2, organoids were treated with 1μM 4-hydroxytamoxifen (4-OH tamoxifen) and the media was replaced the next day with organoid medium supplemented with Rosell 1:100 dilutions of NTBF supernatants, ETBF supernatants or ETBF supernatants containing CDCA (50µM) and after 3-5 additional days in culture, the percentage of organoids with lineage tracing events was quantified. Quantification of lineage tracing and Lgr5-GFP cells was analyzed using Rosell a Nikon Eclipse Ts2 fluorescence microscope with a 20x objective.

### Reactive oxygen species analysis

Intracellular ROS levels in organoids from 1-month AIN76A and NWD1 fed mice were assessed using CellROX™ Green Reagent (Thermo Fisher Scientific, C10444). At 48 hours post-seeding, organoids were incubated with 5μM CellROX™ Green Reagent in organoid culture medium for 30min at 37°C, 5% CO₂. Following incubation, the medium was replaced with fresh organoid culture medium to remove unbound dye. Organoids were imaged daily from days 3-7 post-seeding (days 1-5 post-staining) using a Nikon Eclipse Ts2 fluorescence microscope. Both brightfield and fluorescence images (488 nm excitation) were acquired for each field of view and merged for analysis. ROS quantification was performed on day 5 post organoid-seeding (day 3 post-staining) images using ImageJ software. Individual organoids were manually outlined using the freehand selection tool, and mean fluorescence intensity was measured within each selected region. All images were acquired using identical exposure settings, and fluorescence intensity measurements were performed using consistent threshold parameters across all experimental conditions.

### FACS quantification of *Lgr5*^+^ ISCs

Intestinal crypts from AIN76A and NWD1-fed *Lgr5*^EGFP-IRES-creERT2^ mice were isolated, single cell digested, and subjected to FACS analysis as described above in the section organoid experiments. *Lgr5*^+^ ISCs were identified and quantified by using previously established gates^53^.

### Quantitative PCR analysis

Intestinal crypts isolated from 1-month diet-fed mice were lysed and total RNA was extracted using the QIAwave RNAmini Kit (Qiagen, #74536) according to the manufacturer’s protocol, with final elution in 15 µL RNase-free water. RNA concentration and purity were assessed by NanoDrop spectrophotometry (A260/280 ratio). Complementary DNA (cDNA) was synthesized from 250-500ng of total RNA using the cDNA Reverse Transcriptase Kit (Thermo Fisher, #k1622) according to the manufacturer’s instructions. Gene expression was quantified by RT-qPCR using iTaq Universal SYBR Green Supermix (Biorad, # 1725121) on a qTOWER³ G thermocycler (Analytik Jena) equipped with the Connect™ Real-Time System. Each 20 µL reaction contained 2ng cDNA template, 10 µL 2× SYBR Green Master Mix, and 10uM of each forward and reverse primer (sequences provided in the Supplementary Data File). Thermal cycling conditions were as follows: initial denaturation at 95°C for 3min, followed by 40 cycles of 95°C for 10s, 60°C for 30s, and 72°C for 30s. A melting curve analysis was performed at the end of each run to confirm amplification specificity. All samples were run in technical duplicate alongside a water control. Data were acquired and analyzed using qPCRsoft 4.1 software (Analytik Jena). Relative gene expression was calculated using the 2^(−ΔΔCt) method, with expression of each target gene normalized to the housekeeping gene GAPDH and presented as fold change relative to control conditions. Statistical analysis was performed in GraphPad Prism using unpaired multiple t-tests.

### SDS-PAGE and western-blot analysis

For SDS-PAGE and western blot of organoids, we isolated crypts from AIN76A fed mice as described above. 3 days after seeding organoids were treated for 24 hrs with medium supplemented NTBF or ETBF supernatants, harvested and lysed directly in 2X Laemmli buffer by mechanical disruption and heating at 96°C for 5 minutes, followed by centrifugation at 10,000×g for 5 minutes to remove debris. Equal volumes of each lysate were separated on 10% acrylamide gels and transferred to nitrocellulose membranes (Amersham) using a wet-transfer system at 350mA for 90 minutes at 4°C. Membranes were blocked for 1 hour at room temperature in 5% BSA in TBST (TBS + 0.1% Tween-20), followed by overnight incubation at 4°C with primary antibodies against non-phosphorylated (active) β-catenin (S33/S37/T41) (D13A1) (Cell Signaling Technologies, 8814S) and YAP (D8HIX) (Cell Signaling Technologies, 14074S), antibodies against β-actin (Cell Signalling Technologies, Cat.no.: 4970S, RRID:AB_2223172) were used as loading controls. After washing, membranes were incubated with HRP-conjugated secondary antibodies (Goat-Anti-Rabbit igG (H + L)-HRP Conjugat, Bio-Rad, Cat.no.: 1706515, RRID:AB_11125142, dilution 1: 4000) for 1 hour at room temperature and developed using enhanced chemiluminescence (ECL, Thermo Fisher Scientific). Signal detection was performed using a Bio-Rad ChemiDoc imaging system.

### Fecal DNA extraction and targeted PCR analysis

For microbiome analysis, individual fecal pellets of experimental mice were collected, pooled and stored at −80°C until processing. DNA extraction was performed using the QIAamp Fast DNA Stool mini-Kit (Qiagen) with modifications to improve yield. Briefly, 180-220mg of fecal material was weighed and 1mL InhibitEX Buffer was added. Samples were vortexed, then incubated at 70°C with shaking at 1,100rpm for 5min. Following additional vortexing, samples were centrifuged at 14,000g for 1min at 4°C. The supernatant (200μL) was transferred to a fresh tube containing 15μL Proteinase K, 200μL Buffer AL, and approximately 200μL acid-washed microceramic beads (1.4 mm diameter). Samples were vortexed and incubated at 75°C with shaking at 1,100 rpm for 10mins. The lysate was transferred to a clean tube, and 400μL of absolute ethanol was added and mixed thoroughly. The entire lysate was applied to a QIAamp spin column and centrifuged at 14,000g for 1min at 4°C. Subsequent steps were performed according to the manufacturer’s protocol. DNA concentration and purity were assessed using a Nanodrop 1000 spectrophotometer (Peqlab). DNA was eluted in 15µl of nuclease-free water. DNA integrity was assessed by electrophoresis on 2% agarose gels prepared in TAE buffer (40 mM Tris base, 20 mM acetic acid, 1 mM EDTA), alongside the GeneRuler 100 bp Plus DNA Ladder. Electrophoresis was performed at 120 V for 60 min, and bands were visualized under UV illumination. PCR amplification was performed using species-specific and universal bacterial primers targeting 16S rRNA, *Bacteroides fragilis spp.*, enterotoxigenic *B. fragilis* (ETBF), *Sutterella spp*., and *Lactobacillus spp*. (sequences provided in the Supplementary Data File). The following cycling conditions were used: 16S rRNA: initial denaturation at 95 °C for 5 min; 30 cycles of 94 °C for 1 min, 55 °C for 1 min, and 72 °C for 90 s; final extension at 72 °C for 7 min; *Bacteroides fragilis*: initial denaturation at 94 °C for 5 min; 40 cycles of 94 °C for 20 s, 50 °C for 20 s, and 72 °C for 30 s; final extension at 72 °C for 5 min; Enterotoxigenic *B. fragilis* (ETBF): initial denaturation at 94 °C for 1 min; 40 cycles of 94 °C for 45 s, 52 °C for 45 s, and 72 °C for 45 s; final extension at 72 °C for 7 min; *Sutterella spp.*: initial denaturation at 95 °C for 15 min; 30 cycles of 94 °C for 1 min, 60 °C for 1 min, and 72 °C for 1 min; final extension at 72 °C for 5 min. *Lactobacillus spp.*: initial denaturation at 96 °C for 5 min; 35 cycles of 94 °C for 45 s, 56 °C for 45 s, and 72 °C for 3.5 min; final extension at 72 °C for 10 min. DNA samples were ran on a 2% agarose gel in TAE (Tris-acetate-EDTA: 40mM Tris base, 20mM acetic acid and 1mM EDTA) buffer, together with the GeneRuler 100 bp Plus DNA Ladder. The gel was left for running in TAE buffer at 120V for 1h. The DNA bands were observed under UV light and images were taken. Uncropped images of agarose gels are provided in the Supplementary Data File. DNA from non-enterotoxigenic B. fragilis (NTBF) and ETBF strains served as negative and positive controls. Band intensities of B. fragilis and ETBF were normalized to the corresponding 16S rRNA universal bacterial primer band intensities. Relative abundance was expressed as fold induction over the mean of the control group (n=3).

### Single nucleus RNA sequencing analysis

Colon tissue from CO₂-euthanized mice (n=3 per condition) was snap-frozen in liquid nitrogen and stored at –80°C. Nuclei were extracted on ice using Nuclei Extraction Buffer (Miltenyi Biotec) and a gentle MACS Dissociator, filtered through 70-μm and 30-μm strainers, and pelleted at 300×g. Magnetic enrichment was performed using Anti-Nucleus MicroBeads and LS columns (Miltenyi Biotec). Nuclei quality and viability were assessed by propidium iodide staining on a Cellometer Auto 2000 (Nexcelom). Single-nucleus multiomics libraries were prepared using the BD Rhapsody single-cell ATAC-seq, mRNA whole-transcriptome analysis (WTA), and Sample Tag workflow (Doc ID 23-24799(01)). Isolated nuclei were multiplex labeled with MS Nucleoporin P62 Sample Tags 1, 5, 7, 8, 9, and 10 (Cat. Nos. 460291, 460293, 460294, 460295, 460296, and 460297). For tagging, up to 1 × 10^6 nuclei per sample were resuspended in modified ATAC-SMK buffer containing RNase inhibitor and DTT, incubated with 2 μL Sample Tag reagent on ice for 30 min, washed twice at 500 × g for 5 min at 4 °C, and resuspended in modified nuclei buffer. A subsample was counted on the BD Rhapsody Scanner after DyeCycle Green staining, enabling the pooling of samples at equal ratios. Tagmentation was then performed with 100,000 Sample Tag-labeled nuclei for 30 min at 37 °C. For single-nucleus capture, BD Rhapsody Enhanced Cell Capture Beads V3 (Cat. No. 667052) were converted to splint beads by annealing Universal ATAC-Seq splint oligo, followed by washing and resuspension in sample buffer. Tagmented nuclei were diluted in modified sample buffer with RNase inhibitor, recounted, and loaded onto a primed BD Rhapsody 8-Lane Cartridge (Cat. No. 666262), followed by loading of splint beads. After cartridge capture and washing, nuclei were lysed on cartridge at 37 °C, beads were retrieved and washed, and the captured material was processed by ligation, reverse transcription, splint-oligo removal, and Exonuclease I treatment. Libraries were generated using the BD Rhapsody Tagmentation and Supplemental Reagents Kit (Cat. No. 571201), and WTA Amplification Kit (Cat. No. 633801). AMPure XP beads (Cat. No. A63880) were used for cleanup. Sample Tag and WTA libraries were generated by random priming and extension, Random Priming Extension (RPE) PCR, and WTA index PCR. Final libraries were quantified using the Qubit dsDNA HS assay and quality was assessed on an Agilent Bioanalyzer. Sequencing was performed by Novogene on an Illumina NovaSeq X 25B Flowcell, aiming for a depth of 70.000 reads per nucleus for the WTA libraries and 600 reads per nucleus for the Sample Tag libraries. Raw data was processed using the BD Rhapsody™ Sequence Analysis Pipeline (Revision: 0).. Computational analysis was performed in Python (v3.13.6) using Scanpy (v1.11.4)^55^. Cells flagged as doublets or multiplets by the BD Rhapsody™ pipeline were excluded. Remaining cells were filtered based on the following criteria: ≥300 total counts, 400–10,000 detected genes, and <20% mitochondrial gene content. Genes detected in fewer than 10 cells were removed. Counts were normalized to 10,000 per cell and log-transformed. Highly variable genes were selected for PCA, followed by Harmony-based batch correction (harmonypy v0.0.10)^56^. Cells were clustered using the Leiden algorithm (leidenalg v0.10.2, resolution=0.11)^57^ and visualized by UMAP^58^. Clusters were annotated manually using known marker genes cross-referenced with PanglaoDB^59^, CellMarker^60^, or the indicated published literature. Cell cycle phase was scored using Seurat-derived gene sets via sc.tl.score_genes_cell_cycle^61^. Epithelial cells were isolated and subjected to a second round of preprocessing, batch correction, and re-clustering (resolution=0.6). DEGs were identified using the Wilcoxon rank-sum test via rank_genes_groups. Differentially expressed genes were identified using Scanpy, and genes were ranked by log₂ fold change between conditions. Pre-ranked GSEA was conducted using the GSEApy package (v1.1.9). Gene sets were obtained from the Molecular Signatures Database (MSigDB) Hallmark collection (version h.all.v2025.1) and the Reactome pathway collection (Reactome_2022). Enrichment scores were normalized to yield Normalized Enrichment Scores (NES), and statistical significance was assessed using false discovery rate (FDR) q-values. Pathways were ranked based on NES and visualized using Matplotlib. Pathway names were reformatted for clarity and further grouped and interpreted according to their biological function. Cell-type-specific responses to dietary intervention were quantified using the Augur algorithm^26^ implemented in PertPy. Analysis was performed on epithelial cells comparing Control and 1-month NWD1 samples using the default Augur workflow (random_state = 42). Higher AUC values indicate greater transcriptional responsiveness to the dietary intervention. All visualizations were generated using Scanpy^55^., Matplotlib (v3.10.5) (http://dx.doi.org/10.1109/MCSE.2007.55), and seaborn (v0.13.2) (https://doi.org/10.21105/joss.03021).

### 16S ribosomal RNA (rRNA) gene amplicon sequencing of mouse stool

Fecal DNA was extracted using the phenol-chloroform method, as previously described^62^. The 16S rRNA gene amplification protocol was adapted from the Earth Microbiome Project^63^. The V4 region of the 16S rRNA gene from extracted DNA was amplified by PCR using the Platinum™ Hot Start PCR Master Mix (Cat #: 13000014, Thermo Fisher Scientific) and then purified by the AMPure XP Beads (Cat #: A63880, Beckman Coulter Life Sciences) to remove free primers and primer dimers. The purified amplicon was quantified using a Qubit dsDNA HS assay (Cat #: Q32854, Thermo Fisher Scientific) and equal amount (by mass concentration) was pooled together. Sequencing of the 16S rRNA V4 library was performed on an Illumina MiSeq instrument (Illumina, San Diego, CA) with 150bp paired-end reads at the Harvard Medical School Biopolymer facility. The demultiplexed raw sequencing data were imported to the QIIME2 environment (version 2020.8)^64^. The low-quality bases were trimmed, and the reads were joined, denoised, and checked for chimeras using DADA2 plug-in prior to taxonomic assignment^65^. Taxonomic assignment of each amplicon sequence variant (ASV) was conducted using a pre-trained Naive Bayes classifier with the SILVA database (version 138.1)^66^. The ASVs feature table was further used for differential abundance analysis using MaAslin2^67^. In this analysis, the differential abundance testing focused on the genus-level due to resolution limitation of 16S rRNA V4 region; low abundant genera (average relative abundance below 0.1%) were not included in the statistical testing. Each genus-level was modeled as a function of diet, genotype, and sex (categorical variable) with caging as a random effect. Genus levels with a FDR-corrected p-value of less than 0.25 are considered significant. α-diversity and β-diversity analyses were performed using the phyloseq R package (v1.48.0)^68^ in R (v4.4.0). Taxonomy abundance data are provided as Supplementary Data.

### Statistical analysis

No statistical method was used to predetermine the sample size for *in vivo* experiments. Tables and graphs for statistical analysis were created using GraphPad Prism 4.03 (GraphPad Software). Statistical significance between two groups was determined by two-tailed Student’s t-test, and that for more than two groups was determined by one-way ANOVA with Tukey’s multiple comparison test. Differential bacterial abundance was assessed using linear mixed-effects models as implemented in MaAsLin2. All of the data in the graphs are shown as mean ± s.d., unless stated otherwise.

### Ethics

The organoid work of this study was conducted in accordance with the current versions of the Declaration of Helsinki and the Medical Association’s professional code of conduct in Baden-Württemberg, Germany. It was approved by the ethics committees of the Universities of Heidelberg (S-733/2022) and Mannheim (2012-293N-MA).

### Data representation

For the presentation of our data, the figures were created using Microsoft Power Point Microsoft® PowerPoint® 2016 MSO (Version 2603 Build 16.0.19822.20086) 32 Bit. Illustrations were created with Biorender (Schmitt, M. (2026) https://BioRender.com/r0duioq).

## Data availability

All data supporting the findings of this study are available from the authors upon reasonable request. Single-nucleus RNA sequencing and 16S ribosomal RNA (rRNA) gene amplicon sequencing datasets will be made publicly available upon publication. Detailed information on diet formulations, oligonucleotide sequences, original agarose gel images, sequencing results for *B. fragilis* strain TT12, taxonomy abundance table and MaAsLin2 results of differentially abundant bacteria are included in the Supplementary Data.

## Supporting information

Taxonomy abundance table

MaAsLin2 results

## Acknowledgments

We thank D. Saur for kindly providing the *c-Kit*^CreERT2^ mice, Richard S. Blumberg for providing the *Defa6*^Cre^ mice and Hans Clevers for providing the p*Lys*^CreERT2^ mice. Funding for grants IIG_2014_1181 and IIG_FULL_2022_015 was obtained from Wereld Kanker Onderzoek Fonds (WKOF) as part of the World Cancer Research Fund International grant programme. This project was additionally supported by the Deutsche Forschungsgemeinschaft (DFG) – Project number 550879831. This work was furthermore supported by NIH grant DK088199.

## Author contributions

S. Silva, P. Procopio, V. Zinina were responsible for the study concept and design; data acquisition, analysis, and interpretation; and manuscript revision. R.R.S.M., S.S., S.B. L.L. L.N. A.S.,M.P.V., F.A., K.P., R.J., V.S., H.G., D.S., R.S.B., J.P., W.S.G., were responsible for data acquisition, analysis, and interpretation or provided resources. R.F. was responsible for the study concept and design; data analysis and interpretation, M.S. was responsible for the study concept and design; data analysis and interpretation manuscript writing and revision; obtaining funding; and study supervision.

## Author information

**Mathijs P. Verhagen**

Present affiliation: Discovery Oncology, Genentech Inc., South San Francisco, CA, USA.

---

**Supplementary Figure S1 (related to Figure 1):**
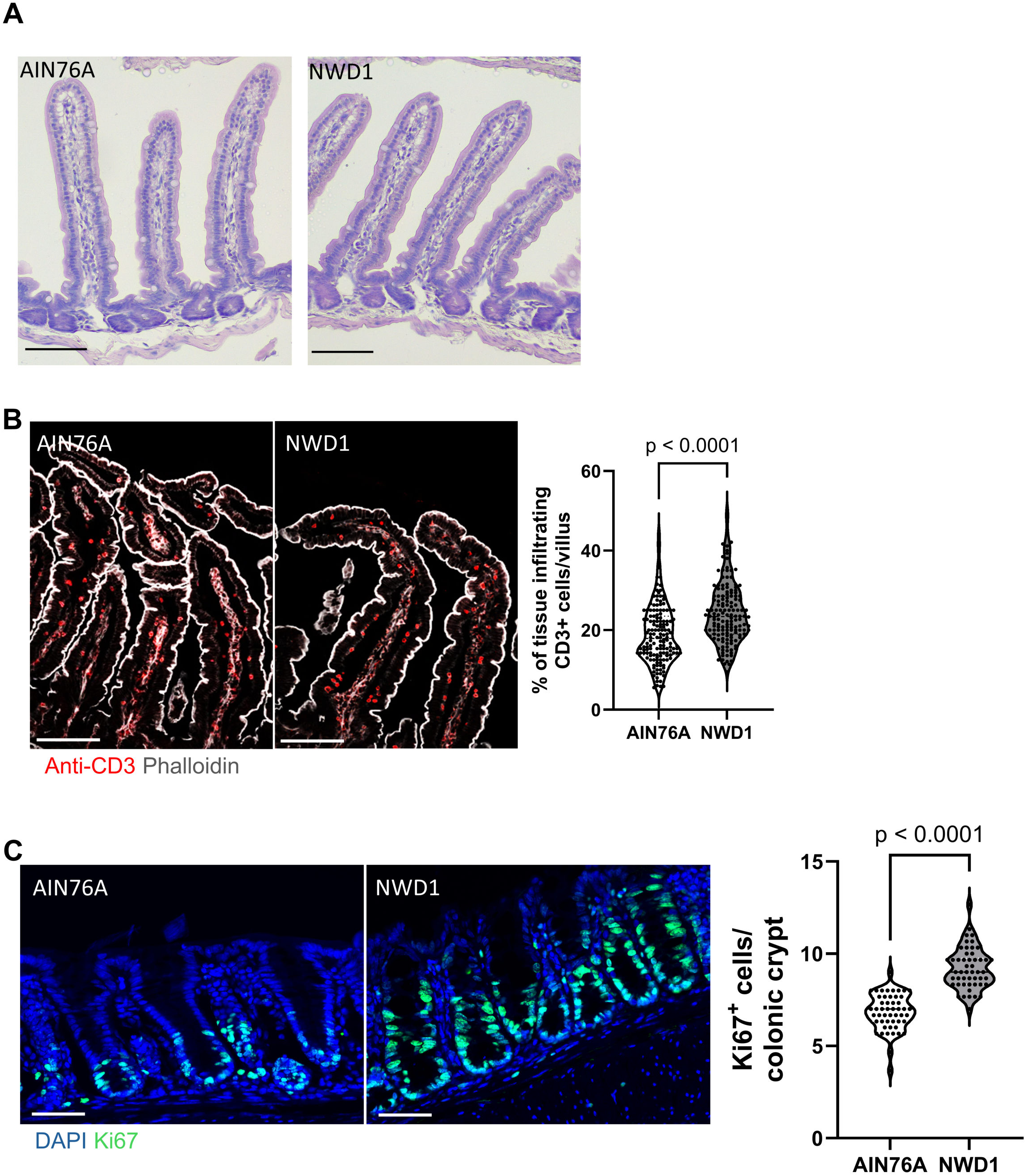
**A)** Hematoxylin and eosin (H&E staining of small intestinal tissues of 1-month diet fed mice (n =3 individual mice per group, scale bar = 75 µm). **B)** Confocal microscopy analysis and quantification of the percentage of CD3^+^ cells infiltrating the small intestinal epithelium of mice fed for 1 month with AIN76A or NWD1 (results represent pooled data of 3 individual mice per group, scale bars = 100 µm, cells were counterstained with phalloidin). **C)** Confocal microscopy analysis and quantification of Ki67^+^ cells in colonic crypts of 1-month AIN76A and NWD1 fed mice (results represent pooled data of 3 individual mice per group, scale bar= 50 µm, nuclei were counterstained with DAPI). All data are mean ± s.d. and were analyzed by two-tailed Student’s t-test (**B** and **C**).

**Supplementary Figure S2 (related to Figure 2):**
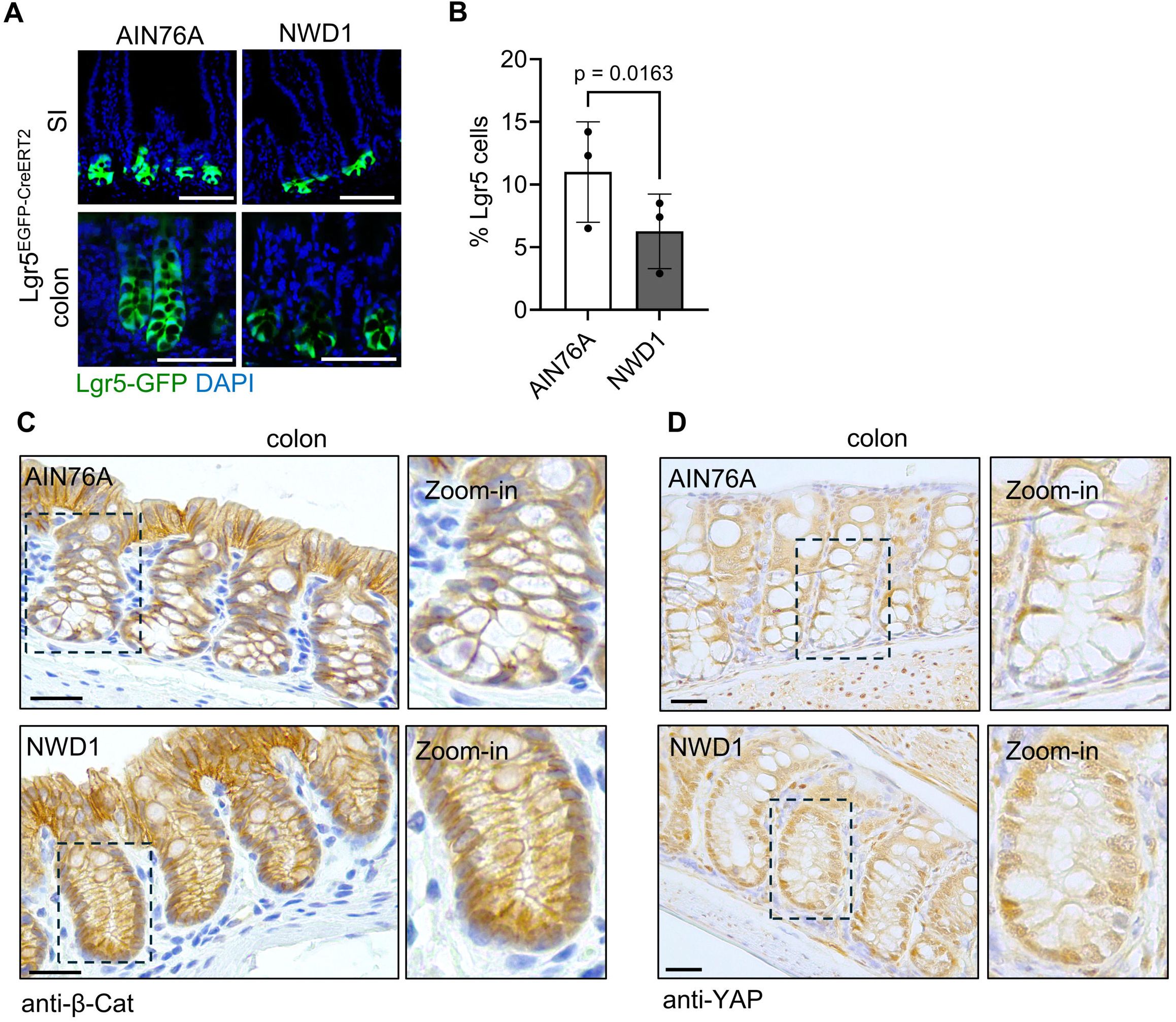
**A)** Confocal microscopy images of intestinal tissues of *Lgr5*^EGFP-IRES-CreERT2^ mice fed with AIN76A or NWD1 for 3 months (n = 3 mice per group; scale bar = 100 µm; nuclei were counterstained with DAPI). **B)** Average percentage of small intestinal *Lgr5*-GFP^+^ cells of AIN76A or NWD1 fed *Lgr5*^EGFP-IRES-CreERT2^ mice, as determined by flow cytometry (n= 3 mice per group). **C)** β-Catenin IHC analysis of colonic tissues of mice fed for 1 month with AIN76A or NWD1 (n=3 per group, scale bar = 100 µm, box marks the area of higher magnification image). **D)** YAP IHC analysis of colonic tissues of mice fed for 1 month with AIN76A or NWD1 (n=3 mice per group, scale bar = 100 µm, box marks the area of higher magnification image). Data shown in (**B**) are mean ± s.d. and were analyzed by or two-tailed Student’s *t*-test.

**Supplementary Figure S3 (related to Figure 3):**
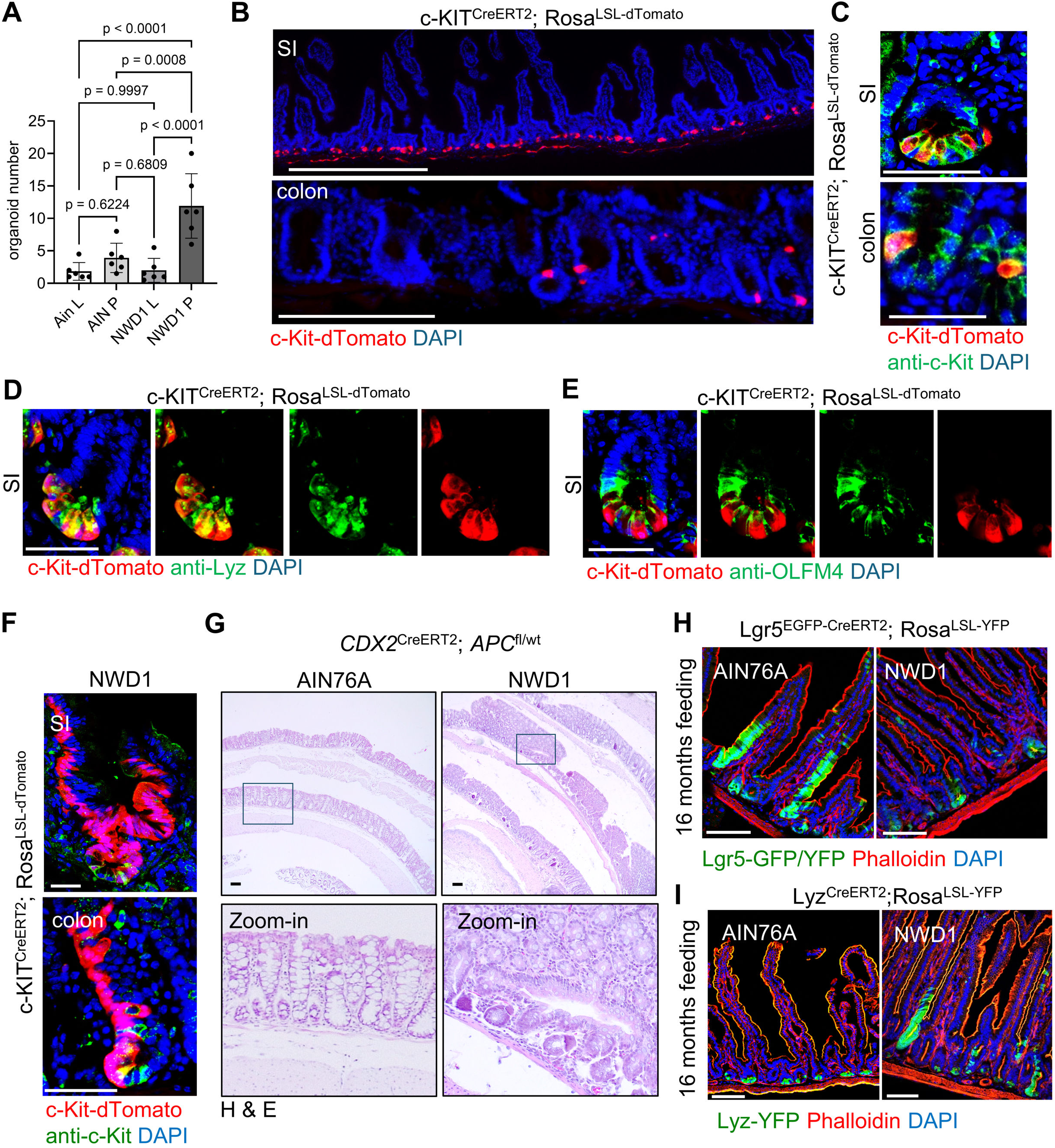
**A)** Organoid multiplicities derived from FACS sorted single *Lgr5*^+^ ISCs or PCs from *Lgr5*^EGFP-IRES-CreERT2^ mice fed for 3 months with AIN76A or NWD1 (L= *Lgr5*^+^ ISCs; P= PCs) (n = 3 independent experiments performed in duplicates). **B)** Confocal microscopy images of intestinal tissues of AIN76A fed *c-Kit*^CreERT2^/*R26*^LSL-dTomato^ mice 10 days after tamoxifen treatment. (n=3, scale bars = 500 µm, nuclei were counterstained with DAPI). **C)** Analysis of c-Kit and dTomato expression in AIN76A fed *c-Kit*^CreERT2^/*R26*^LSL-dTomato^ mice 10 days after tamoxifen treatment (n=3, scale bars = 50 µm, nuclei were counterstained with DAPI). **D)** Analysis of lysozyme expression in AIN76A fed *c-Kit*^CreERT2^/*R26*^LSL-dTomato^ mice 10 days after Tamoxifen treatment (n=3, scale bars = 50 µm, nuclei were counterstained with DAPI). **E)** Analysis of OLFM4 expression in AIN76A fed *c-Kit*^CreERT2^/*R26*^LSL-dTomato^ mice 10 days after Tamoxifen treatment (n=3, scale bars = 50 µm, nuclei were counterstained with DAPI). **F)** Analysis of c-Kit and dTomato expression in NWD1 fed *c-Kit*^CreERT2^/*R26*^LSL-dTomato^ mice 10 days after tamoxifen injection, showing lineage tracings originating from *c-Kit*^+^ cells located at the SI and colonic crypt base (n=3, nuclei were counterstained with DAPI). **G)** H&E staining of colonic tissues from *CDX2*^CreERT2^; *APC*^fl/wt^ mice fed for 6 months with AIN76A or NWD1 diets after tamoxifen induction (n = 5 mice per group, scale bar = 100 µm). Box indicates region of hoghermagnification images. **H** and **I)** Confocal microscopy analysis of *Lgr5*^EGFP-IRES-creERT2^/*R26*^LSL-YFP^ (H) or *Lys*^CreERT2^/*R26*^LSL-YFP^ (I) mice, fed with AIN76A or NWD1 diets for 16 months (n=2, scale bar = 100 µm, tissues were counterstained with Phalloidin and DAPI). Data shown in (**A**) are mean ± s.d. and were analyzed by one-way ANOVA with Tuckey’s multiple comparison test

**Supplementary Figure S4 (related to Figure 4):**
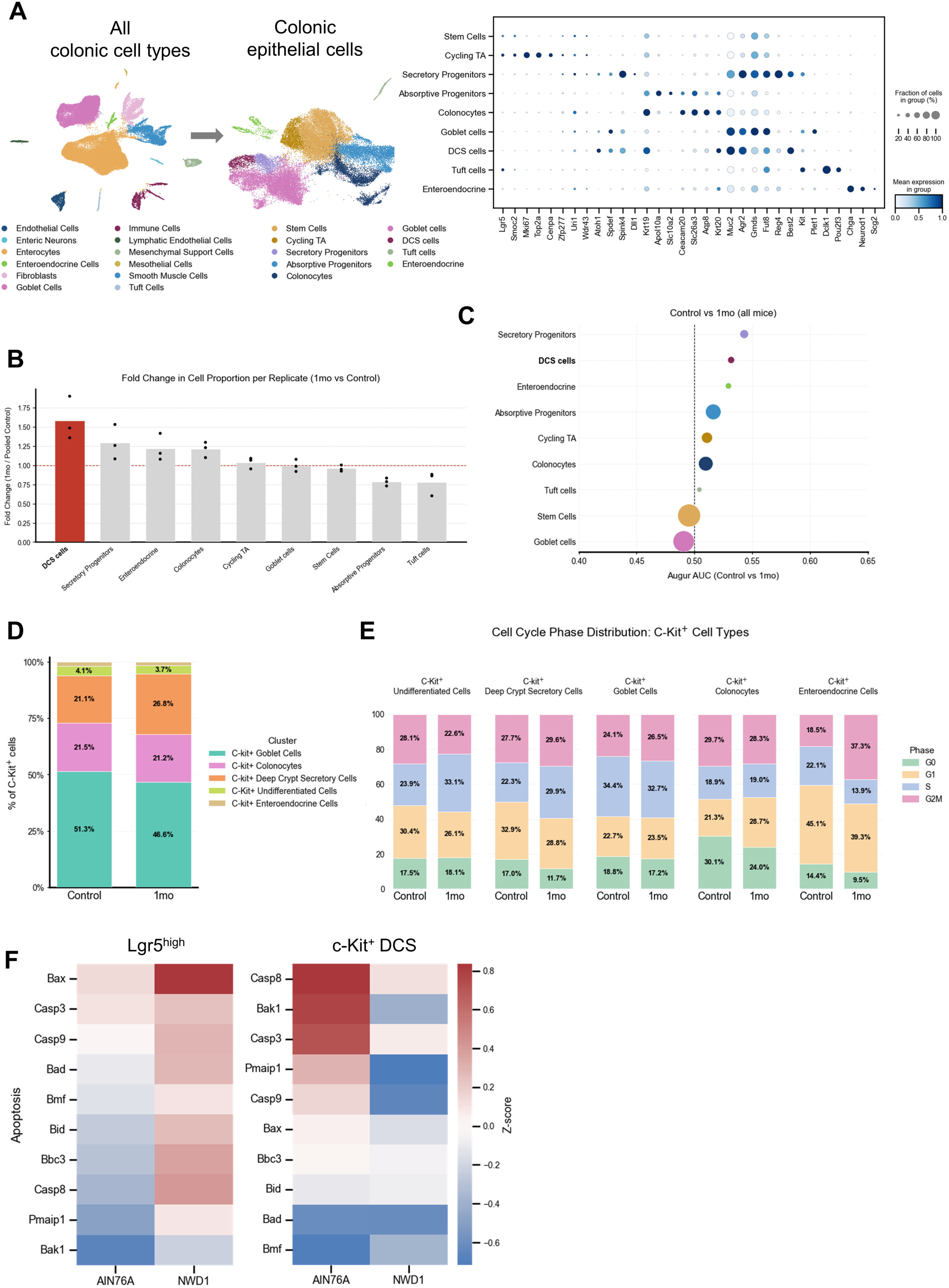
**A)** UMAP embedding of all intestinal cell types (left) and intestinal epithelial cell clusters (right), annotated according to the depicted canonical marker gene expression (n = 3 mice per condition). **B)** Bar plot showing the fold change in cellular abundance of epithelial lineages defined in (A), comparing NWD1-fed mice at 1 month to pooled AIN76A controls. Bar height represents the mean fold change across replicates, with individual data points showing per-mouse values. The dashed line indicates no change (fold change = 1). DCS cells are highlighted (n = 3 mice per condition). **C)** Augur dot plot of cell type prioritization. Visualization of cell-type-specific responses to 1 month of NWD1. Dot size indicates the percentage of cells expressing the gene/feature within each cluster. Dot color reflects the average expression level. Cell types are ordered on the y-axis by their overall Augur cell type prioritization score (n = 3 mice per condition). **D)** Distribution of cellular lineages across c-Kit^+^ intestinal epithelial cell types of mice fed for 1 month with AIN76A or NWD1 (n=3 mice per group. **E)** Bar plots of cell cycle phase distribution across *c-Kit*^+^ cell subclusters as defined in Fig. 3A in 1-month AIN76A and NWD1-fed mice. Cell cycle phases were scored using Seurat cell cycle gene signatures (Scanpy implementation) (n = 3 mice per condition). **F)** Heatmap depicting z-scored expression of apoptosis-related genes in *Lgr5*^high^ ISCs (left) and *c-Kit*^+^ DCS cells (right) comparing 1-week AIN76A and NWD1-fed mice (n = 3 individual mice per condition).

**Supplementary Figure S5 (related to Figure 5):**
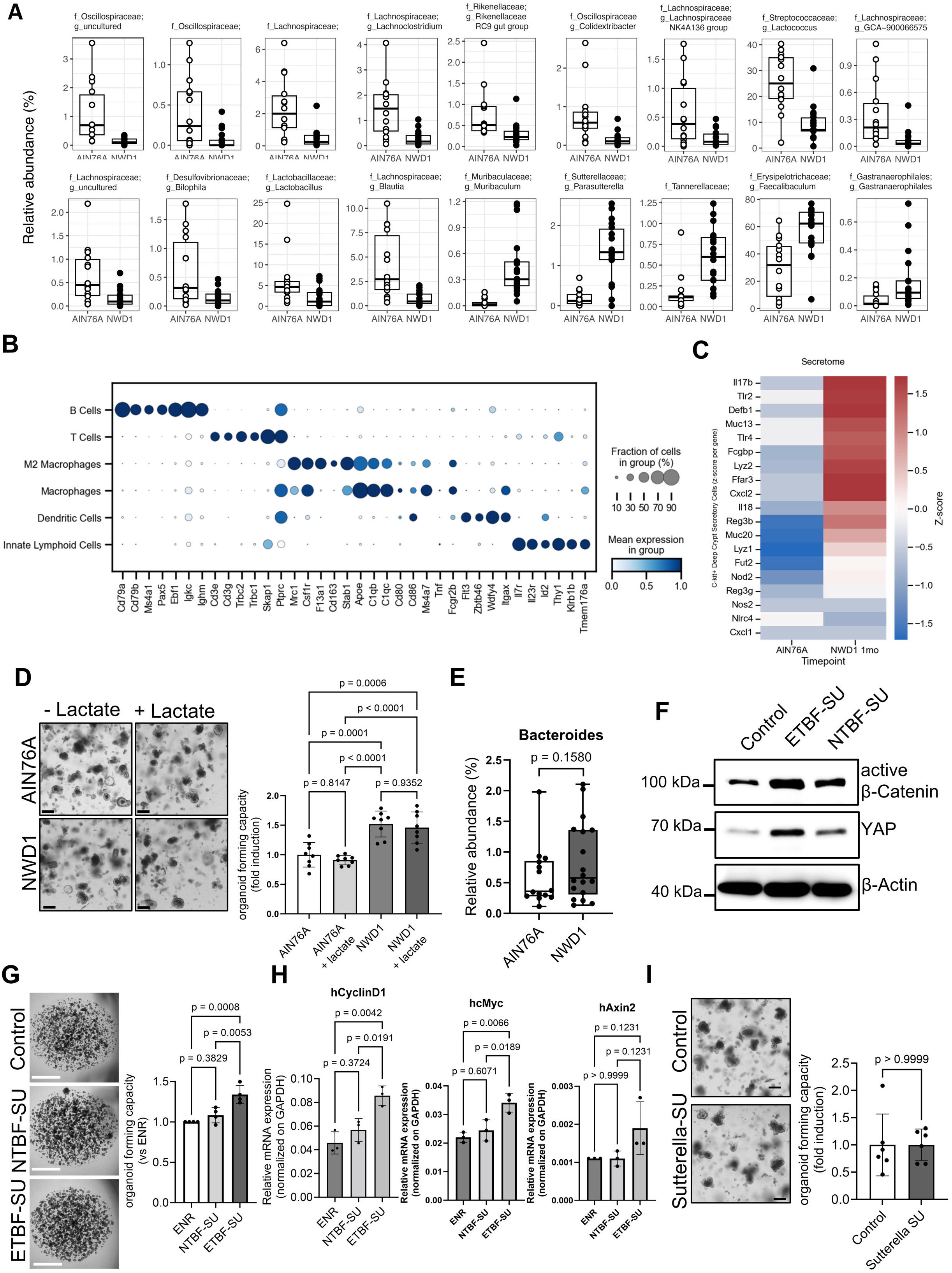
**A)** Boxplot indicating the relative abundance of the indicated bacterial genera in 3-months AIN76A and NWD1 fed mice according to 16S rRNA gene amplicon sequencing analysis (n = 14 for AIN76A and 18 for NWD1). Statistical analysis was performed using a linear mixed-effects model implemented in MaAsLin2. **B)** Transcriptional profiles used to cluster immune cell types in the UMAP plot shown in Fig. 5D (n= 3 mice per condition). **C)** Heatmap depicting z-scored expression of secretome genes in *c-Kit*^+^ DCS cells comparing 1-month AIN76A and NWD1-fed mice (n = 3 individual mice per condition). **D)** Organoid multiplicities of SI crypts of mice fed with AIN76A or NWD1 for 1 month, cultured in the presence or absence of lactate (n = 4 biological replicates, scale bar = 200 µm). **E)** Boxplot indicating the relative abundance of *Bacteroides* in 3-months AIN76A and NWD1 fed mice according to 16S rRNA gene amplicon sequencing analysis (n = 14 for AIN76A and 18 for NWD1). Statistical analysis was performed using a linear mixed-effects model implemented in MaAsLin2. **F)** Western blot analysis of YAP and β-Catenin protein levels in SI organoids treated with NTBF-SU or NTBF-SU. One representative out of 4 individual experiments is shown. Antibodies against β-actin were used as loading control. **G)** Multiplicities of human organoids cultured in organoid medium, or organoid medium supplemented with NTBF or ETBF supernatants (n=2 experiments performed in duplicates). **H)** RT–qPCR analysis of Wnt-target genes in organoids cultured in organoid medium, or organoid medium supplemented with NTBF or ETBF supernatants (n=1 experiments performed in triplicates). **I)** Organoid multiplicities of murine SI crypts cultured in medium supplemented with *Sutterella wadsworthensis* supernatant (Sutterella-SU) or the corresponding bacterial growth medium (control) (n = 3 biological replicates performed in duplicates, scale bar = 200 µm). All data are mean ± s.d. and were analyzed using two-tailed student’s t-test (**I**) or one-way ANOVA (**D**, **G**, **H**) with Tuckey’s multiple comparison test.

**Supplementary Figure S6 (related to Figure 6):**
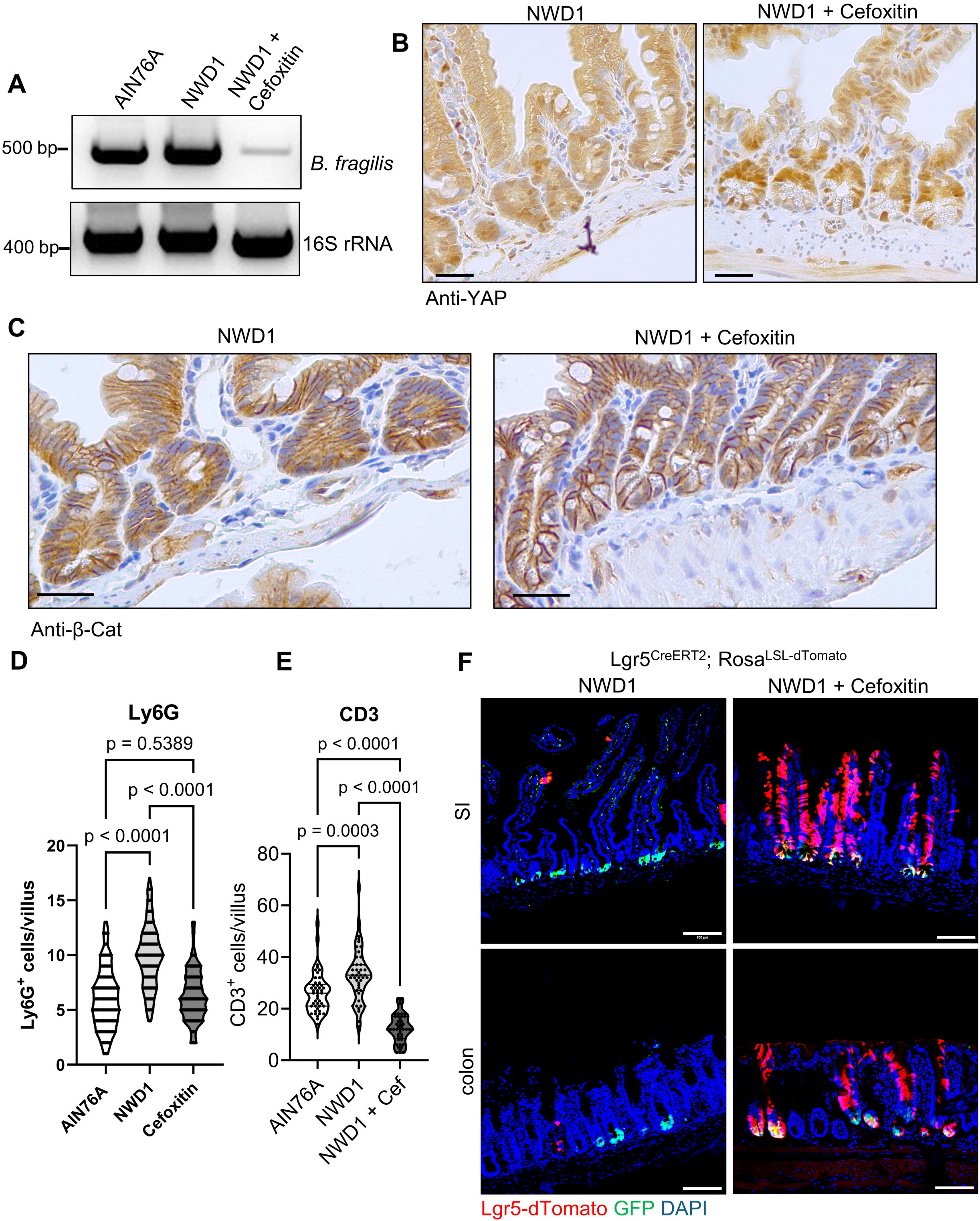
**A)** PCR analysis of stool DNA from mice fed with NWD1 in the presence or absence of Cefoxitin. *B. fragilis* DNA was used positive control for *B. fragilis* primers. 16s primers detecting all bacteria were used as loading control (n=3 biological replicates). **B)** YAP IHC analysis of mouse small intestinal tissues from NWD1 fed mice simultaneously treated with Cefoxitin (n=3 mice per group, scale bar = 50 µm). **C)** β-Catenin IHC of mouse small intestinal tissues from NWD1 fed mice simultaneously treated with Cefoxitin (n=3 mice per group, scale bar = 50 µm). **D and E)** Analysis of Ly6G (D) and CD3 (E) expression in mice fed with AIN76A or NWD1 diets in the presence or absence of Cefoxitin (scale bars = 500 µm, nuclei were counterstained with DAPI). Graph shows the quantification of total CD3^+^ cell numbers/villus (results represent pooled data of 3 individual mice per group). **F)** Confocal microscopy images of intestinal tissues of *Lgr5*^EGFP-IRES-CreERT2^/*Rosa*^LSL-dTomato^ mice that received NWD1 diets for 1 month in the presence or absence of Cefoxitin (n = 5 mice per group, scale bar = 100 µm, nuclei were counterstained with DAPI). All data are mean ± s.d. and were analyzed by one-way ANOVA with Tuckey’s multiple comparison test.

