## Supplementary Data File for "Western-style diet-induced microbial shifts and inflammation reprogram *c-Kit*⁺ secretory cells into targets for tumor initiation"

*to whom correspondence should be addressed at:

Dr. Mark Schmitt

#### Table of content:

**Supplementary Data Figure 1:** Blast report of enterotoxigenic *B. fragilis* strain T12, used for the inoculation of mice used in *in vivo* experiments described in Figure 6F

**Supplementary Data Figure 2:** Uncropped agarose gel images of PCR results shown in Figure 5C

**Supplementary Data Figure 3:** Uncropped agarose gel images of PCR results shown in Figure 5G

**Supplementary Data Figure 4:** Uncropped agarose gel images of PCR results shown in Supplementary Figure S6A

**Supplementary Data Figure 5:** Uncropped gel images of immunoblot results shown in Supplementary Figure S5F.

**Supplementary Table 1**: Diet formulation of AIN76A diets used in this study

**Supplementary Table 2**: Diet formulation of NWD1 diets used in this study

**Supplementary Table 3**: List of primers used in this study

### Supplementary Data Figure 1

Blast report of enterotoxigenic *B. fragilis* strain T12

**Bacteroides fragilis bft-1 gene for metalloprotease, complete cds**

Sequence ID: AB026625.1 Length: 1546 Number of Matches: 1

Range 1: 989 to 1322 GenBank Graphics ▼ Next Match ▲ Previous Match

| Score | Expect | Identities | Gaps | Strand |
|---|---|---|---|---|
| 590 bits(319) | 6e-164 | 329/334(99%) | 0/334(0%) | Plus/Minus |

```
Query  1     CCGTATTCACATTATAAGAATTGAACCAGGACATCCCTAAGATTTTATTATCCCAAGTAC  60
             ||||||||||||||||||||||||||||||||||||||||||||||||||||||||||||
Sbjct  1322  CCGTATTCACATTATAAGAATTGAACCAGGACATCCCTAAGATTTTATTATCCCAAGTAC  1263

Query  61    CCCAGCGTATTAAAAAATAAATTTGATCGTCATAACCTTCTGCTTTTGGATTACTTTTTA  120
             ||||||||||||||||||||||||||||||||||||||||||||||||||||||||||||
Sbjct  1262  CCCAGCGTATTAAAAAATAAATTTGATCGTCATAACCTTCTGCTTTTGGATTACTTTTTA  1203

Query  121   GTGAAGCAGTAAAGCCTTCCAGTCCCTCTTTGGCGTCGCCACTTGGACAACTGTATTCAG  180
             ||||||||||||||||||||||||||||||||||||||||||||||||||||||||||||
Sbjct  1202  GTGAAGCAGTAAAGCCTTCCAGTCCCTCTTTGGCGTCGCCACTTGGACAACTGTATTCAG  1143

Query  181   TAGTATACAGTACAAAGTGGAAATTGACATATCTTTTCAGTCCATGAACTGCATAAACCG  240
             ||||||||||||||||||||||||||||||||||||||||||||||||||||||||||||
Sbjct  1142  TAGTATACAGTACAAAGTGGAAATTGACATATCTTTTCAGTCCATGAACTGCATAAACCG  1083

Query  241   AGTTCGCCGCATCCTGCATCTGGGCACTAACTTCATTAGGATAGATAGTACTTCCATTCT  300
             ||||||||||||||||||||||||||||||||||||||||||||||||||||||||||||
Sbjct  1082  AGTTCGCCGCATCCTGCATCTGGGCACTAACTTCATTAGGATAGATAGTACTTCCATTCT  1023

Query  301   CTCTCAGACAAATCACATACACCGTCTTCGGCTC  334
             ||||||||||||| ||||| ||||| || || ||
Sbjct  1022  CTCTCAGACAAATGACATATACCGTTTTAGGTTC  989
```

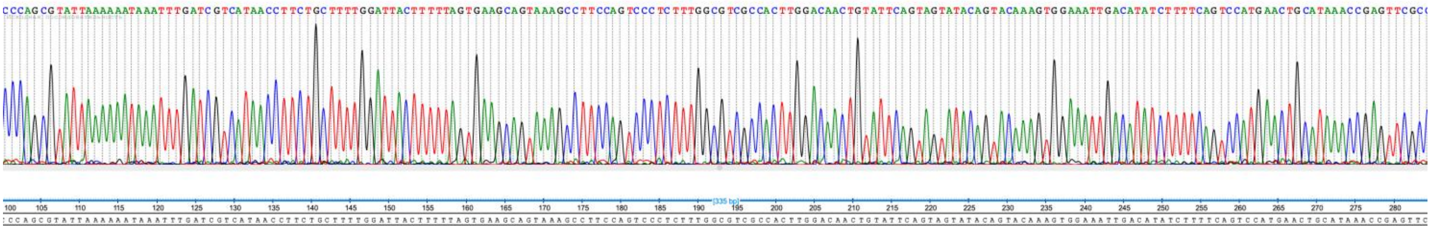

**Supplementary Data Figure 2**

Uncropped agarose gel images of PCR results shown in Figure 5C. Only relevant lanes and bands are labeled. Red boxes indicate how the gel was cropped for the final figure.

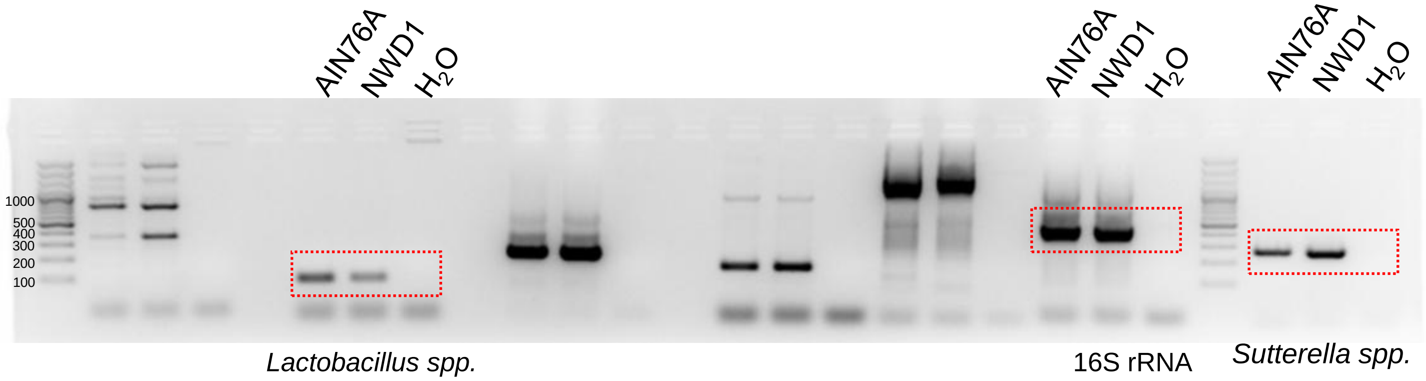

**Supplementary Data Figure 3**

Uncropped agarose gel images of PCR results shown in Figure 5G. Only relevant lanes and bands are labeled. Red boxes indicate how the gel was cropped for the final figure.

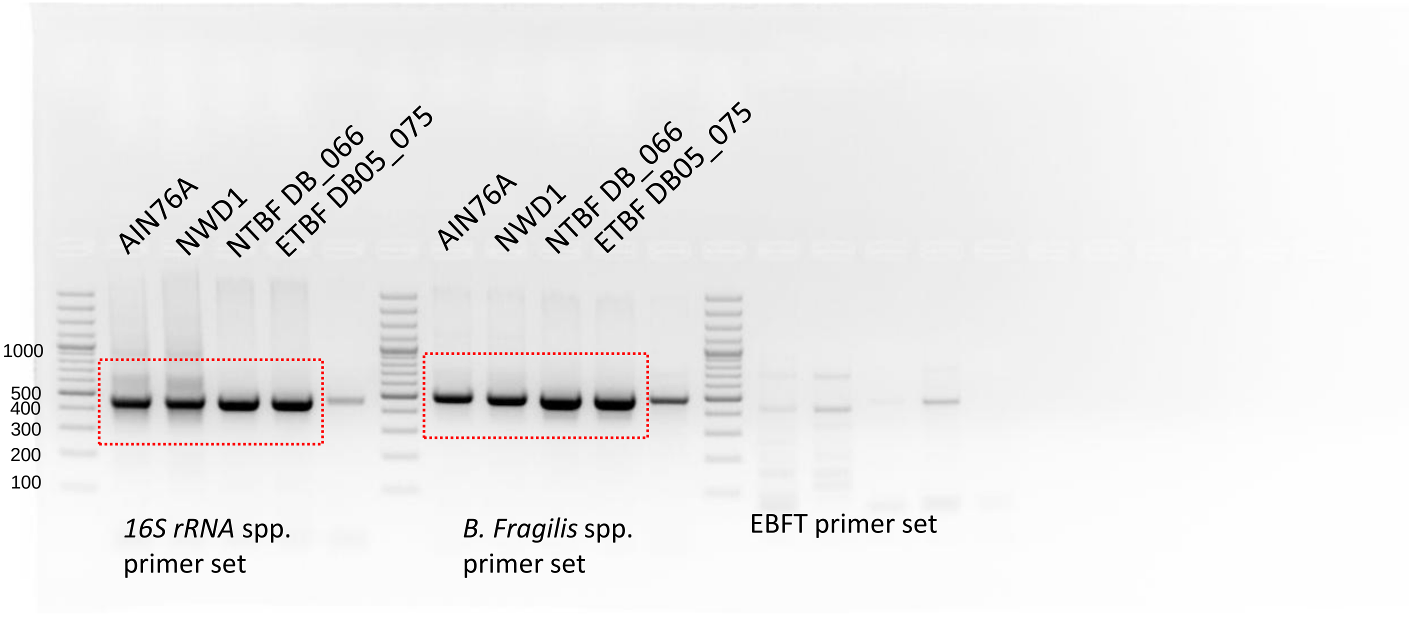


Higher exposure

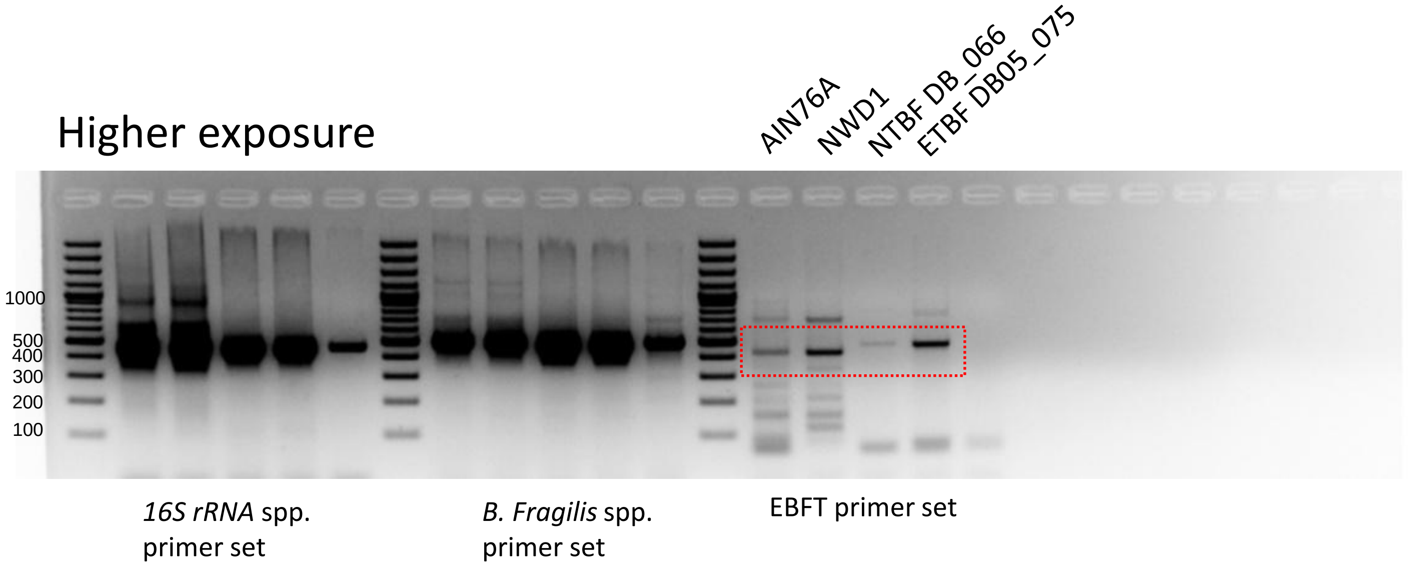

**Supplementary Data Figure 4**

Uncropped agarose gel images of PCR results shown in Supplementary Figure S6A. Only relevant lanes and bands are labeled. Red boxes indicate how the gel was cropped for the final figure.

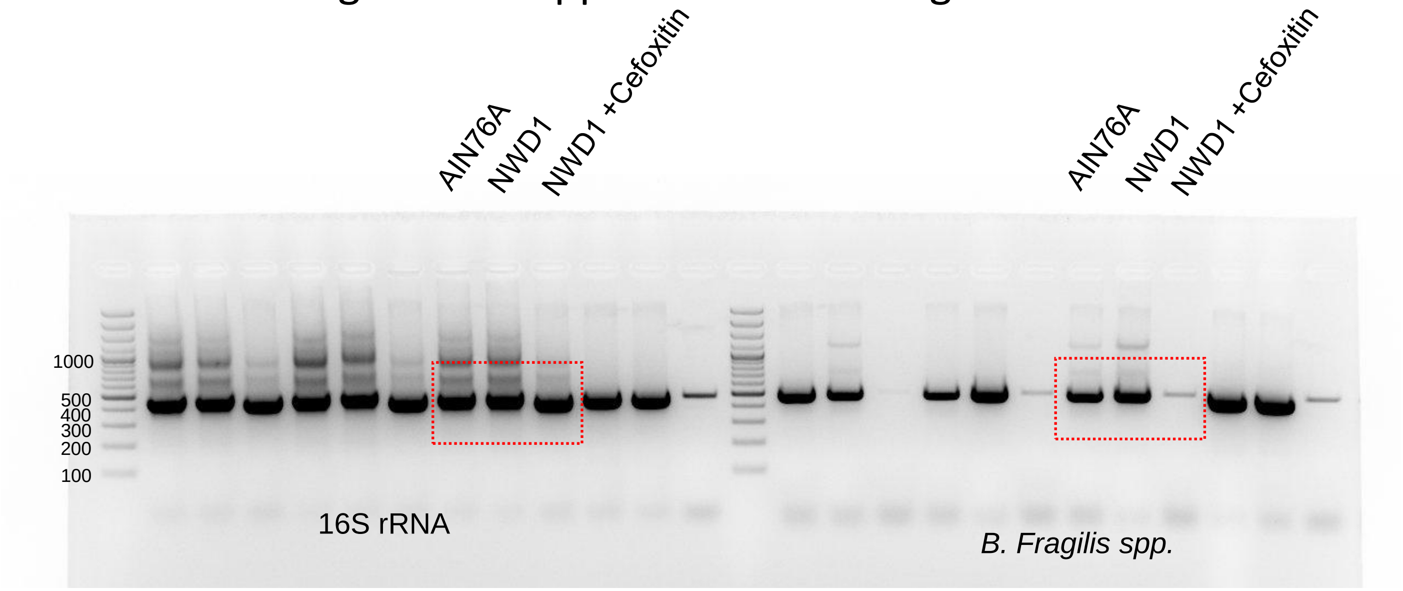

**Supplementary Data Figure 5**

Uncropped gel images of immunoblot results shown in Supplementary Figure S5F. Only relevant lanes and bands are labeled. Red boxes indicate how the gel was cropped for the final figure.

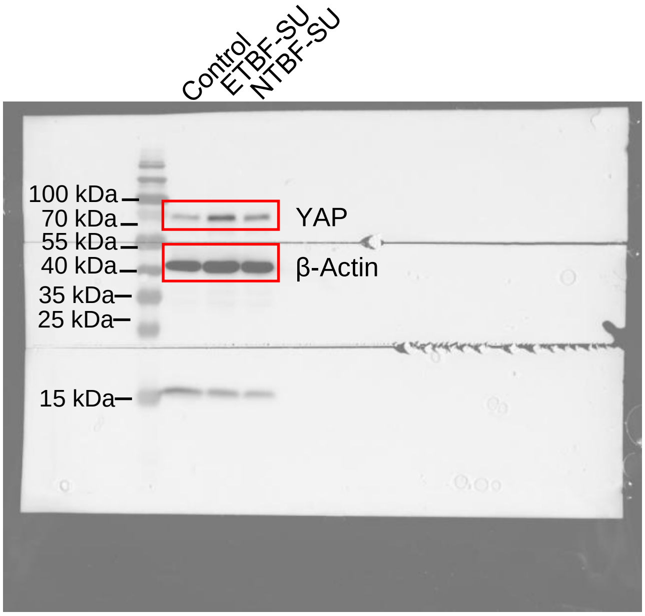


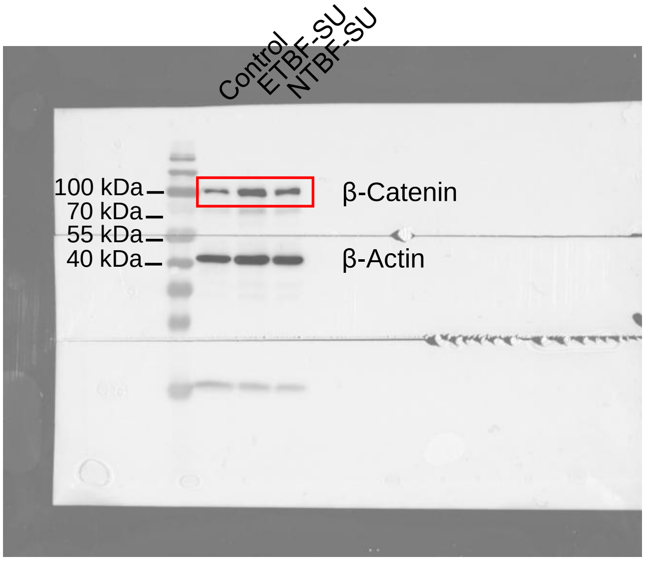

**Supplementary Table 1**

Diet formulation of AIN76A diets used in this study

Altromin Spezialfutter GmbH & Co. KG
Im Seelenkamp 20 - D-32791 Lage

**25.4.2023 13:34 Seite: 1**

**Nummer** **Lfd. Nummer** Änderung 25.04.2023 13:33:48
6043 1 AIN - 76 A Purified Diet, Rats or Mouse Research Diets D10001

| Nr. | Inhaltsstoff | Einheit | Bedarf | Gehalt | Differenz |
|---|---|---|---|---|---|
| 1 | Rohprotein / Crude Protein | mg/kg | | 206157,688 | |
| 2 | Rohfett / Crude Fat | mg/kg | | 53842,212 | |
| 3 | Rohfaser / Crude Fibre | mg/kg | | 38054,321 | |
| 4 | Rohasche / Crude Ash | mg/kg | | 36351,846 | |
| 5 | Feuchtigkeit / Moisture | mg/kg | | 68392,355 | |
| 6 | Monosaccharide(s) | mg/kg | | 11477,525 | |
| 7 | Disaccharide(s) | mg/kg | | 157726,500 | |
| 8 | Polysaccharide(s) | mg/kg | | 401720,750 | |
| 9 | Umsetzb. Energie/Metab. Energy | kcal/kg | | 3634,486 | |
| 10 | Lysin / Lysine | mg/kg | | 20844,523 | |
| 11 | Methionin / Methionine | mg/kg | | 11414,162 | |
| 12 | Cystin / Cystine | mg/kg | | 3807,605 | |
| 13 | Threonin / Threonine | mg/kg | | 8535,692 | |
| 14 | Tryptophan | mg/kg | | 2364,148 | |
| 15 | Arginin / Arginine | mg/kg | | 11731,267 | |
| 16 | Histidin / Histidine | mg/kg | | 6293,612 | |
| 17 | Isoleucin / Isoleucine | mg/kg | | 8616,745 | |
| 18 | Leucin / Leucine | mg/kg | | 17554,856 | |
| 19 | Phenylalanin / Phenylalanine | mg/kg | | 8540,970 | |
| 20 | Valin / Valine | mg/kg | | 3892,186 | |
| 21 | Alanin / Alanine | mg/kg | | 2926,840 | |
| 22 | Asparaginsäure / Aspartic acid | mg/kg | | 4209,831 | |
| 23 | Glutaminsäure / Glutamic acid | mg/kg | | 28160,035 | |
| 24 | Glycin / Glycine | mg/kg | | 3710,176 | |
| 25 | Prolin / Proline | mg/kg | | 15193,066 | |
| 26 | Serin / Serine | mg/kg | | 6252,655 | |
| 27 | Tyrosin / Tyrosine | mg/kg | | 11085,598 | |
| 28 | Vitamin A | I.E./kg | | 4050,000 | |
| 29 | Vitamin D3 | I.E./kg | | 1050,000 | |
| 30 | Vitamin E | mg/kg | | 88,611 | |
| 31 | Vitamin K3 als/as Menadion(e) | mg/kg | | 0,500 | |
| 32 | Vitamin B1 | mg/kg | | 50,048 | |
| 33 | Vitamin B2 | mg/kg | | 60,386 | |
| 34 | Vitamin B6 | mg/kg | | 60,041 | |
| 35 | Vitamin B12 | mg/kg | | 0,263 | |
| 36 | Nikotinsäure / Nicotinic acid | mg/kg | | 300,204 | |
| 37 | Pantothensre./Pantothenic acid | mg/kg | | 150,127 | |
| 38 | Folsäure / Folic acid | mg/kg | | 2,00288 | |
| 39 | Biotin | mg/kg | | 0,201 | |
| 40 | Cholinchlorid/Choline chloride | mg/kg | | 1013,802 | |
| 43 | Inosit / Inositol | mg/kg | | 13,202 | |
| 45 | Calcium | mg/kg | | 4347,009 | |
| 46 | Ges.Phosphor / Phosphorus | mg/kg | | 5186,986 | |
| 47 | Verd.Phosphor/Digest.Phosporus | mg/kg | | 5016,961 | |
| 48 | Magnesium | mg/kg | | 565,910 | |
| 49 | Natrium / Sodium | mg/kg | | 2079,615 | |
| 50 | Kalium / Potassium | mg/kg | | 3782,660 | |
| 51 | Schwefel / Sulfur | mg/kg | | 2131,595 | |
| 52 | Chlor / Chlorine | mg/kg | | 1499,440 | |
| 53 | Eisen / Iron | mg/kg | | 45,954 | |

**Seite: 2**

| Nr. | Inhaltsstoff | Einheit | Bedarf | Gehalt | Differenz |
|---|---|---|---|---|---|
| 54 | Mangan / Manganese | mg/kg | | 51,402 | |
| 55 | Zink / Zinc | mg/kg | | 40,556 | |
| 56 | Kupfer / Copper | mg/kg | | 6,751 | |
| 57 | Jod / Iodine | mg/kg | | 0,229 | |
| 59 | Fluor / Fluorine | mg/kg | | 4,000 | |
| 60 | Selen / Selenium | mg/kg | | 0,219 | |
| 61 | Kobalt / Cobalt | mg/kg | | 0,018 | |
| 63 | Caprinsäure C-10:0 | mg/kg | | 53,000 | |
| 64 | Laurinsäure C-12:0 | mg/kg | | 53,000 | |
| 65 | Myristinsäue C-14:0 | mg/kg | | 53,000 | |
| 66 | Pentadecansäure C-15:0 | mg/kg | | 53,000 | |
| 67 | Palmitinsäure C-16:0 | mg/kg | | 6466,000 | |
| 68 | Palmitoleinsäure C-16:1 | mg/kg | | 53,000 | |
| 69 | Margarinsäure | mg/kg | | 53,000 | |
| 70 | Stearinsäure C-18:0 | mg/kg | | 1272,000 | |
| 71 | Ölsäure C-18:1 | mg/kg | | 12243,000 | |
| 72 | Linolsäure C-18:2 | mg/kg | | 28991,000 | |
| 73 | Linolensäure C-18:3 | mg/kg | | 1431,000 | |
| 74 | Arachinsäure C-20:0 | mg/kg | | 477,000 | |
| 75 | Eicosaensäure C-20:1 | mg/kg | | 636,000 | |
| 76 | Eicosadiensäure C-20:2 | mg/kg | | 477,000 | |
| 77 | Arachidonsäure C-20:4 | mg/kg | | 53,000 | |
| 78 | Eicosapentaensäure C-20:5 | mg/kg | | 53,000 | |
| 79 | Behensäure C-22:0 | mg/kg | | 318,000 | |
| 80 | Essigsäure C- 2:0 | mg/kg | | 53,000 | |
| 81 | Docosahexaensäure C-22:6 | mg/kg | | 53,000 | |
| 82 | Tricosansäure | mg/kg | | 53,000 | |
| 83 | Nervonsäure C-24:1 | mg/kg | | 53,000 | |
| 110 | Chrom / Chromium | mg/kg | | 2,060 | |
| 111 | Aluminium | mg/kg | | 3,404 | |
| 155 | Volumen / Volume | g | | 1000,000 | |

### Supplementary Table 2

Diet formulation of NWD1 diets used in this study

Altromin Spezialfutter GmbH & Co. KG
Im Seelenkamp 20 - D-32791 Lage



**Nummer** **Lfd. Nummer** Änderung 25.04.2023 11:24:52

130008 1 Newmark Stress Diet C ( D16378C) basierend auf AIN- 76A

| Nr. | Inhaltsstoff | Einheit | Bedarf | Gehalt | Differen |
|---|---|---|---|---|---|
| 1 | Rohprotein / Crude Protein | mg/kg | | 206494,959 | |
| 2 | Rohfett / Crude Fat | mg/kg | | 208571,958 | |
| 3 | Rohfaser / Crude Fibre | mg/kg | | 19000,828 | |
| 4 | Rohasche / Crude Ash | mg/kg | | 21958,497 | |
| 5 | Feuchtigkeit / Moisture | mg/kg | | 44310,197 | |
| 6 | Monosaccharide(s) | mg/kg | | 13525,875 | |
| 7 | Disaccharide(s) | mg/kg | | 304399,939 | |
| 8 | Polysaccharide(s) | mg/kg | | 162009,968 | |
| 9 | Umsetzb. Energie/Metab. Energy | kcal/kg | | 4669,680 | |
| 10 | Lysin / Lysine | mg/kg | | 20975,857 | |
| 11 | Methionin / Methionine | mg/kg | | 11458,598 | |
| 12 | Cystin / Cystine | mg/kg | | 3810,953 | |
| 13 | Threonin / Threonine | mg/kg | | 8555,939 | |
| 14 | Tryptophan | mg/kg | | 2375,088 | |
| 15 | Arginin / Arginine | mg/kg | | 11763,485 | |
| 16 | Histidin / Histidine | mg/kg | | 6307,586 | |
| 17 | Isoleucin / Isoleucine | mg/kg | | 8636,344 | |
| 18 | Leucin / Leucine | mg/kg | | 17538,777 | |
| 19 | Phenylalanin / Phenylalanine | mg/kg | | 8545,539 | |
| 20 | Valin / Valine | mg/kg | | 3861,641 | |
| 21 | Alanin / Alanine | mg/kg | | 2845,959 | |
| 22 | Asparaginsäure / Aspartic acid | mg/kg | | 4155,681 | |
| 23 | Glutaminsäure / Glutamic acid | mg/kg | | 28141,555 | |
| 24 | Glycin / Glycine | mg/kg | | 3688,099 | |
| 25 | Prolin / Proline | mg/kg | | 15195,151 | |
| 26 | Serin / Serine | mg/kg | | 6235,639 | |
| 27 | Tyrosin / Tyrosine | mg/kg | | 11119,351 | |
| 28 | Vitamin A | I.E./kg | | 4049,999 | |
| 29 | Vitamin D3 | I.E./kg | | 120,000 | |
| 30 | Vitamin E | mg/kg | | 89,800 | |
| 31 | Vitamin K3 als/as Menadion(e) | mg/kg | | 0,500 | |
| 32 | Vitamin B1 | mg/kg | | 50,048 | |
| 33 | Vitamin B2 | mg/kg | | 60,390 | |
| 34 | Vitamin B6 | mg/kg | | 60,041 | |
| 35 | Vitamin B12 | mg/kg | | 0,264 | |
| 36 | Nikotinsäure / Nicotinic acid | mg/kg | | 300,206 | |
| 37 | Pantothensre./Pantothenic acid | mg/kg | | 150,128 | |
| 38 | Folsäure / Folic acid | mg/kg | | 1,20290 | |
| 39 | Biotin | mg/kg | | 0,201 | |
| 40 | Cholinchlorid/Choline chloride | mg/kg | | 1013,915 | |
| 43 | Inosit / Inositol | mg/kg | | 13,310 | |
| 45 | Calcium | mg/kg | | 390,302 | |
| 46 | Ges.Phosphor / Phosphorus | mg/kg | | 3795,844 | |
| 47 | Verd.Phosphor/Digest.Phosporus | mg/kg | | 3749,088 | |
| 48 | Magnesium | mg/kg | | 563,609 | |
| 49 | Natrium / Sodium | mg/kg | | 2086,999 | |
| 50 | Kalium / Potassium | mg/kg | | 3781,968 | |
| 51 | Schwefel / Sulfur | mg/kg | | 2102,567 | |
| 52 | Chlor / Chlorine | mg/kg | | 1565,772 | |
| 53 | Eisen / Iron | mg/kg | | 45,717 | |



| Nr. | Inhaltsstoff | Einheit | Bedarf | Gehalt | Differenz |
|---|---|---|---|---|---|
| 54 | Mangan / Manganese | mg/kg | | 51,847 | |
| 55 | Zink / Zinc | mg/kg | | 40,554 | |
| 56 | Kupfer / Copper | mg/kg | | 6,835 | |
| 57 | Jod / Iodine | mg/kg | | 0,103 | |
| 59 | Fluor / Fluorine | mg/kg | | 0,042 | |
| 60 | Selen / Selenium | mg/kg | | 0,211 | |
| 61 | Kobalt / Cobalt | mg/kg | | 0,008 | |
| 63 | Caprinsäure C-10:0 | mg/kg | | 208,000 | |
| 64 | Laurinsäure C-12:0 | mg/kg | | 208,000 | |
| 65 | Myristinsäue C-14:0 | mg/kg | | 208,000 | |
| 66 | Pentadecansäure C-15:0 | mg/kg | | 208,000 | |
| 67 | Palmitinsäure C-16:0 | mg/kg | | 25375,995 | |
| 68 | Palmitoleinsäure C-16:1 | mg/kg | | 208,000 | |
| 69 | Margarinsäure | mg/kg | | 208,000 | |
| 70 | Stearinsäure C-18:0 | mg/kg | | 4991,999 | |
| 71 | Ölsäure C-18:1 | mg/kg | | 48047,990 | |
| 72 | Linolsäure C-18:2 | mg/kg | | 113775,977 | |
| 73 | Linolensäure C-18:3 | mg/kg | | 5615,999 | |
| 74 | Arachinsäure C-20:0 | mg/kg | | 1872,000 | |
| 75 | Eicosaensäure C-20:1 | mg/kg | | 2496,000 | |
| 76 | Eicosadiensäure C-20:2 | mg/kg | | 1872,000 | |
| 77 | Arachidonsäure C-20:4 | mg/kg | | 208,000 | |
| 78 | Eicosapentaensäure C-20:5 | mg/kg | | 208,000 | |
| 79 | Behensäure C-22:0 | mg/kg | | 1248,000 | |
| 80 | Essigsäure C- 2:0 | mg/kg | | 208,000 | |
| 81 | Docosahexaensäure C-22:6 | mg/kg | | 208,000 | |
| 82 | Tricosansäure | mg/kg | | 208,000 | |
| 83 | Nervonsäure C-24:1 | mg/kg | | 208,000 | |
| 110 | Chrom / Chromium | mg/kg | | 2,060 | |
| 111 | Aluminium | mg/kg | | 2,082 | |
| 155 | Volumen / Volume | kg | | 1000,000 | |

**Supplementary Table 3**

List of primers used in this study

| Primers | Forward | Reverse |
|---|---|---|
| mGAPDH | AGACGGCCGCATCTTCTTGT | TGCCGTTGAATTTGCCGTGA |
| mAscl2 | TGGTTAGGGGGCTACTGAGC | CACTTGGCATTTGGTCAGGCT |
| mClu | GATGATCCACCAGGCTCAACAG | ACACAGTGCGGTCATCTTCACC |
| mCD44 | CCACAGCCTCCTTTCAATAACC | GGAGTCTTCGCTTGGGGTA |
| mEphB2 | AGGGGTTGTCACCAAGAGCA | AGGTCACGGTGCACGTAGTT |
| mC-Myc | TCGCTGCTGTCCTCCGAGTCC | GGTTTGCCTCTTCTCCACAGAC |
| mCycD2 | GCAGAAGGACATCCAACCGTAC | ACTCCAGCCAAGAAACGGTCCA |
| mAxin2 | GAGATGACGCCTGTGGAACC | CCTGCTCAGACCCCTCCTTT |
| mAreg | GCAGATACATCGAGAACCTGGAG | CCTTGTCATCCTCGCTGTGAGT |
| mEreg | CAGGCAGTTATCAGCACAACCG | CATGCAAGCAGTAGCCGTCCAT |
| mIL-1α | ACGGCTGAGTTTCAGTGAGACC | CACTCTGGTAGGTGTAAGGTGC |
| mIL-1β | GGGTCCGTCAACTTCAAAGA | TGAAGCAGCTATGGCAACTG |
| mIL-10 | CGGGAAGACAATAACTGCACCC | CGGTTAGCAGTATGTTGTCCAGC |
| mIFNg | CAGCAACAGCAAGGCGAAAAAGG | TTTCCGCTTCCTGAGGCTGGAT |
| mTGFβ1 | TGATACGCCTGAGTGGCTGTCT | CACAAGAGCAGTGAGCGCTGAA |
| mTNFα | CAGGAGGGAGAACAGAAACTCCA | CCTGGTTGGCTGCTTGCTT |
| hAxin2 | CAAACTTTCGCCAACCGTGGTTG | GGTGCAAAGACATAGCCAGAACC |
| hcMyc | CCTGGTGCTCCATGAGGAGAC | CAGACTCTGACCTTTTGCCAGG |
| hCyclinD1 | TCTACACCGACAACTCCATCCG | TCTGGCATTTTGGAGAGGAAGTG |
| hGAPDH | AGCCACATCGCTCAGACAC | GCCCAATACGACCAAATCC |
| 16S rRNA | TCCTACGGGAGGCAGCAGT | GGACTACCAGGGTATCTAATCCTGTT |
| *B. fragilis* | ATAGCCTTTCGAAAGRAAGAT | CCAGTATCAACTGCAATTTTA |
| Enterotoxigenic *B. fragilis* | GAGCCGAAGACGGTGTATGTGATTTGT | TGCTCAGCGCCCAGTATATGACCTAGT |
| *Sutterella spp.* | CGCGAAAAACCTTACCTAGCC | GACGTGTGAGGCCCTAGCC |
| *Lactobacillus spp.* | TGGATGCCTTGGCACTAGGA | AAATCTCCGGATCAAAGCTTACTTAT |
